# Standardized genome-wide function prediction enables comparative functional genomics: a new application area for Gene Ontologies in plants

**DOI:** 10.1101/2021.04.25.441366

**Authors:** Leila Fattel, Dennis Psaroudakis, Colleen F. Yanarella, Kevin O. Chiteri, Haley A. Dostalik, Parnal Joshi, Dollye C. Starr, Ha Vu, Kokulapalan Wimalanathan, Carolyn J. Lawrence-Dill

**Affiliations:** Department of Agronomy, Iowa State University; Department of Plant Pathology and Microbiology, Iowa State University; Department of Ecology, Evolution and Organismal Biology, Iowa State University; Department of Veterinary Microbiology and Preventative Medicine, Iowa State University; Department of Genetics, Development and Cell Biology, Iowa State University

**Keywords:** Gene function, ontology, plants, comparative genomics, functional genomics

## Abstract

**Background:** Genome-wide gene function annotations are useful for hypothesis generation and for prioritizing candidate genes potentially responsible for phenotypes of interest. We functionally annotated the genes of 18 crop plant genomes across 14 species using the GOMAP pipeline.

**Results:** By comparison to existing GO annotation datasets, GOMAP-generated datasets cover more genes, contain more GO terms, and produce datasets similar in quality (based on precision and recall metrics using existing gold standards as the basis for comparison). From there, we sought to determine whether the datasets across multiple species could be used together to carry out comparative functional genomics analyses in plants. To test the idea and as a proof of concept, we created dendrograms of functional relatedness based on terms assigned for all 18 genomes. These dendrograms were compared to well-established species-level evolutionary phylogenies to determine whether trees derived were in agreement with known evolutionary relationships, which they largely are. Where discrepancies were observed, we determined branch support based on jack-knifing then removed individual annotation sets by genome to identify the annotation sets causing unexpected relationships.

**Conclusions:** GOMAP-derived functional annotations used together across multiple species generally retain sufficient biological signal to recover known phylogenetic relationships based on genome-wide functional similarities, indicating that comparative functional genomics across species based on GO data hold promise for generating novel hypotheses about comparative gene function and traits.

## I. BACKGROUND

Phenotypes and traits have long been the primary inspiration for biological investigation. Phenotypes are the result of a complex interplay between functions of genes and environmental cues. In an effort to organize and model gene functions, various systems of classification have been developed including systems like KEGG (the Kyoto Encyclopedia of Genes and Genomes), which is focused on protein function including gene activities superimposed on metabolic pathways [1]. Other such systems include the various Cyc databases, MapMan, and the Gene Ontologies (GO), a vocabulary of gene functions organized as a directed acyclic graph, which makes it innately tractable for computational analysis [2, 3, 4]. GO-based gene function annotation involves the association of GO terms to individual genes. Functions may be assigned to genes based on different types of evidence for the association. For example, functional predictions can be inferred from experiments (EXP), expression patterns (IEP), and more [5]. Computational pipelines are often used to generate functional predictions for newly sequenced genomes, where the genome is first sequenced and assembled, then gene structures (gene models) are predicted, then functions are associated with those gene predictions. Genome-wide gene function prediction datasets are frequently used to analyze gene expression studies, to prioritize candidate genes linked to a phenotype of interest, to design experiments aimed at characterizing functions of genes, and more [6, 7, 8]. How well a gene function prediction set models reality is influenced by how complete and correct the underlying genome assembly and gene structure annotations are as well as by how well the software used to predict functions performs.

GOMAP (the Gene Ontology Meta Annotator for Plants) is a gene function prediction pipeline for plants that generates high-coverage and reproducible functional annotations [9]. The system employs multiple functional prediction approaches, including sequence similarity, protein domain presence, and mixed-method pipelines developed to compete in the Critical Assessment of Function Annotation (CAFA) Challenge [10], a community challenge that has advanced the performance of gene function prediction pipelines over the course of five organized competitions [11].

We previously annotated gene functions for the maize B73 genome and demonstrated that GOMAP’s predicted functions were closer to curated gene-term associations from the literature than those of other community functional annotation datasets, including those produced by Gramene (Ensembl pipeline) and Phytozome (Interpro2GO pipeline) [12]. Using the newly containerized GOMAP system [9], we report here the functional annotation of 18 plant genomes across the 14 crop plant species shown in Table I and report comparisons of performance based on comparison to a Gold Standard gene function datasets, where possible.

**Table I:** Functional annotation sets generated by GOMAP. More information about each dataset including the source of the input to GOMAP can be found at the respective DOI.

| Species | Germplasm/Line | Assembly/Annotation | Dataset DOI | Genome Reference |
| --- | --- | --- | --- | --- |
| *Arachis hypogaea* | Tifrunner | Arachis hypogaea assembly 1.0 | [14] | [15] |
| *Brachypodium distachyon* | Bd21 | Bd21.v3.1.r1 | [16] | [17] |
| *Cannabis sativa* | Hemp | NCBI Cannabis sativa GCA_900626175.1 | [18] | [19] |
| *Glycine max* | Williams 82 | Joint Genome Institute (JGI) Wm82.a4.v1 | [20] | [21] |
| *Gossypium raimondii* | Cotton D | Gossypium raimondii JGI v2.1 | [22] | [23] |
| *Hordeum vulgare* | — | IBSC.PGSB.r1 | [24] | [25] |
| *Medicago truncatula* | R108_HM340 | R108: v1.0 | [26] | [27] |
| *Medicago truncatula* | A17_HM341 | Mt4.0v2 | [28] | [29] |
| *Oryza sativa* | Japonica | IRGSP 1.0 | [30] | [31] |
| *Phaseolus vulgaris* | G19833 | DOE-JGI and USDA-NIFA annotation 2.0 | [32] | [33] |
| *Pinus lambertiana* | Sugar Pine | TreeGenesDB sugar pine assembly v1.5 | [34] | [35] |
| *Sorghum bicolor* | BTx623 | BTx623.v3.0.1.r1 | [36] | [37] |
| *Triticum aestivum* | Chinese Spring | IWGSC RefSeq 1.1 | [38] | [39] |
| *Vigna unguiculata* | IT97K-499-35 | JGI annotation v1.1 | [40] | [41] |
| *Zea mays* * | Mo17 | Zm-Mo17-REFERENCE-CAU-1.0 | [42] | [43] |
| *Zea mays* * | PH207 | Zm-PH207-REFERENCE_NS-UIUC-UMN-1.0 | [44] | [45] |
| *Zea mays* * | W22 | Zm-W22-REFERENCE-NRGene-2.0 Zm00004b.1 | [46] | [47] |
| *Zea mays* * | B73 | RefGen.V4 Zm00001d.2 | [48] | [49] |

Latest overview at https://dill-picl.org/projects/gomap/gomap-datasets/
\* Previously published in [9].

Given these multiple annotations across various plant species, we next considered whether these datasets could be used together for comparative functional genomics in plants. We describe here a simple and crude method by which we used gene function annotations to generate dendrograms of genome-level similarity in function. This idea is similar to that of Zhu et al., who determined the evolutionary relationships among microorganisms based on whole-genome functional similarity [13]. Here we expand on that approach, analyzing genome-wide GO assignments to generate parsimony and distance-based dendrograms (see Figure 1 for process overview). We compared these with well-established species phylogenies (Figure 2) to determine whether trees derived from gene function show any agreement with evolutionary histories, taking agreement between generated dendrograms and known evolutionary histories to be evidence that sufficient comparative biological signal exists to begin to use GO functional annotations across multiple plant genomes for comparative functional genomics investigations.

**Figure 1:**
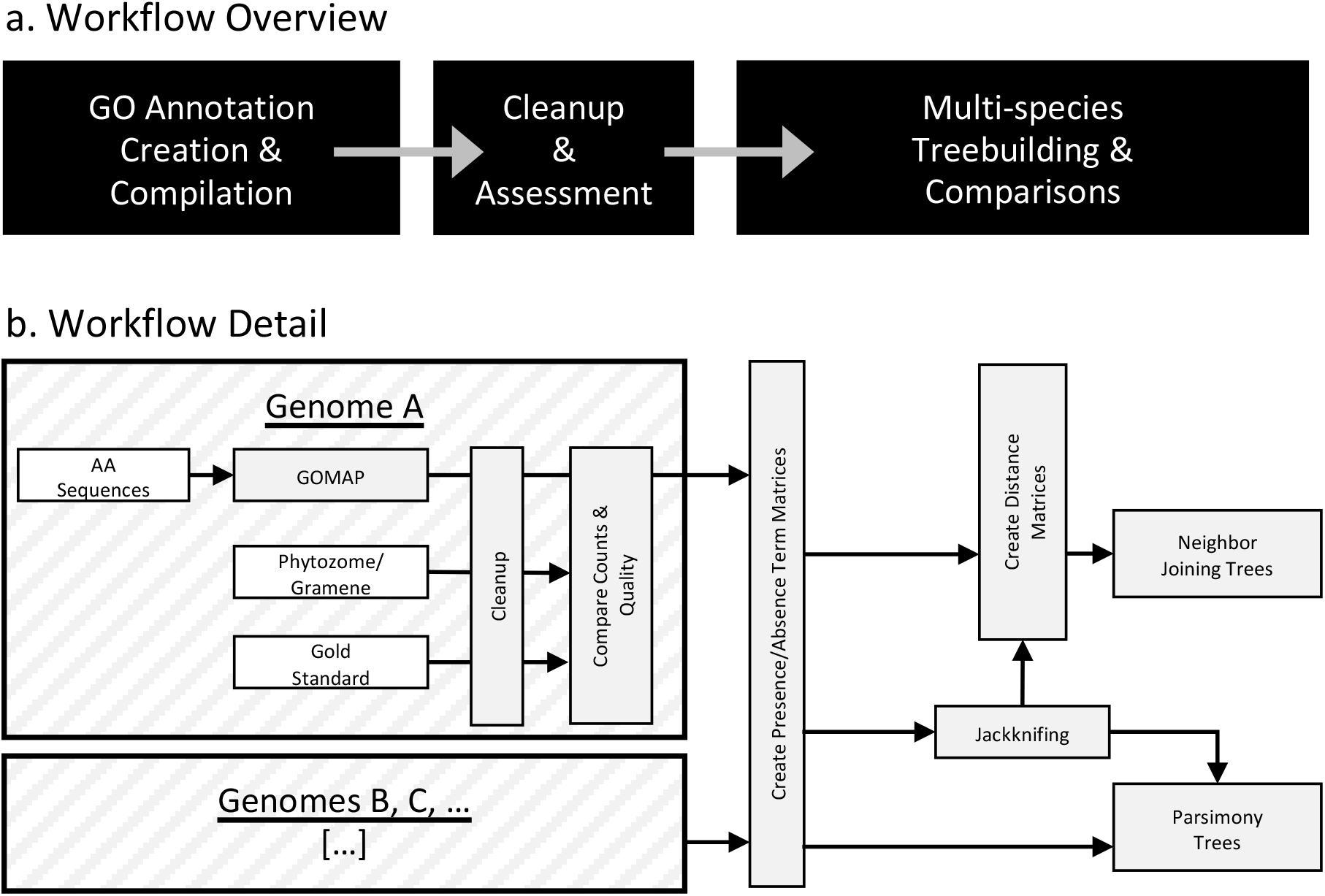
Data workflow schema. The workflow overview is shown in panel ‘a’ with steps represented as black boxes and the flow of information and processes indicated by arrows. Details are shown in panel ‘b’ where the upper large hatched box shows process detail for a single genome and the lower hatched box represents additional genomes for which the details of processing are identical. White boxes represent input datasets. Arrows indicate the flow of information and processes.

**Figure 2:**
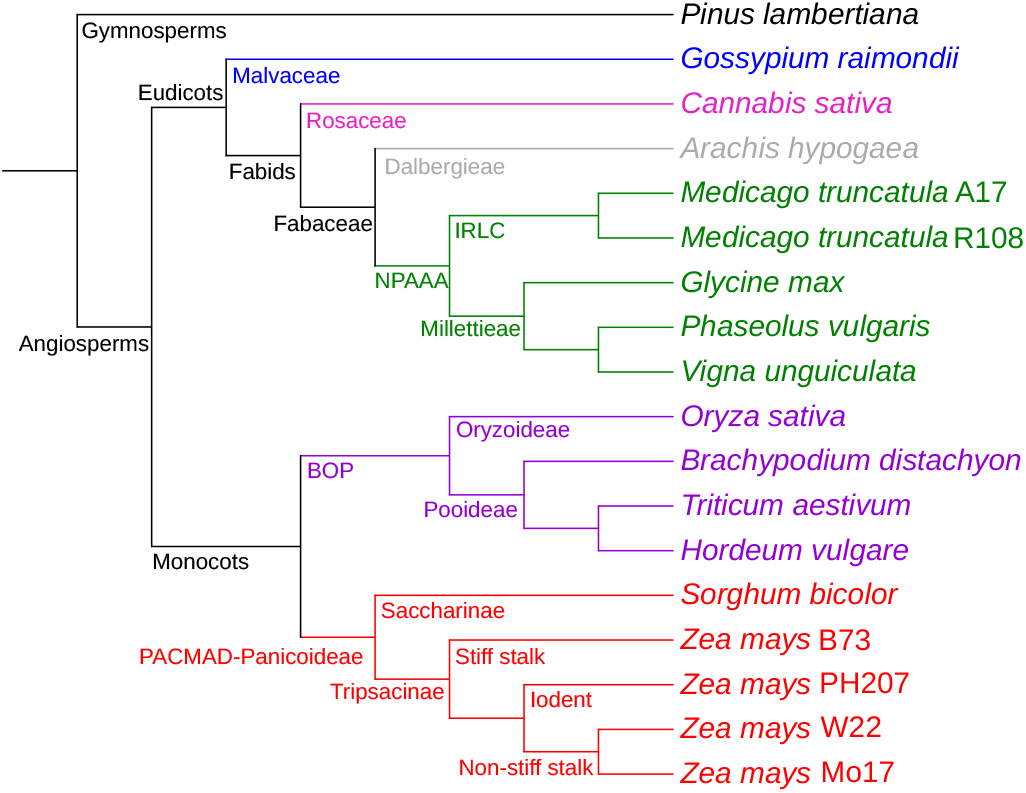
Known phylogenetic relationships among species. Cladogram is rooted by the gymnosperm *Pinus lambertiana* (black). Among angiosperms, eudicots clades include Malvaceae (blue), Rosaceae (magenta), Dalbergieae (grey), and NPAAA (green). Monocots include members of the BOP (purple) and PACMAD-Panicoideae (red) clades.

## II. RESULTS OF ANALYSES

### A. Overview

As shown in Figure 1, gene function annotation sets were created and compiled for each genome. For those with existing annotation sets available on Gramene or Phytozome [50, 51], the datasets were compared. From there, matrices that included genomes as rows and terms as columns were generated. These were used directly to build parsimony trees or to create distance matrices for neighbor-joining tree construction [52, 53, 54]. In subsequent analyses, jackknifing was used to remove terms (columns) or to remove genomes (rows) to map the source of signal for treebuilding results [55].

### B. Functional Annotation Sets Produced

Table II shows quantitative attributes of each of the annotation sets. In summary, GOMAP covers all annotated genomes with at least one annotation per gene, and provides between 3.8 and 12.1 times as many annotations as Gramene or Phytozome.

**Table II:**
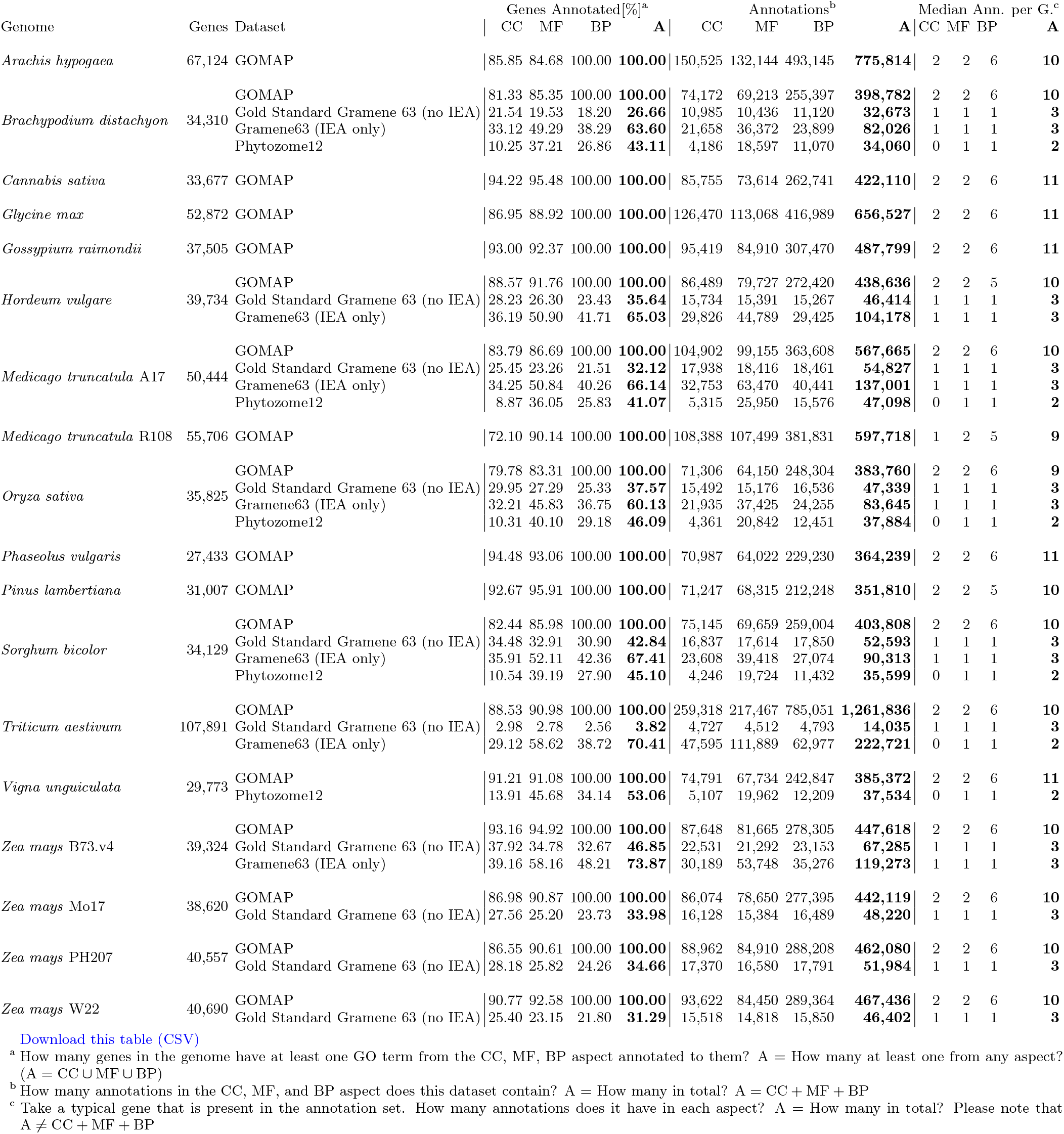
Quantitative metrics of the cleaned functional annotation sets. CC, MF, BP, and A refer to the aspects of the Gene Ontology: Cellular Component, Molecular Function, Biological Process, and Any/All. GOMAP covers all genomes with at least one annotation per gene and provides substantially more annotations than Gramene63 or Phytozome, especially in the BP aspect. The total number of annotations per dataset is visualized in Figure S1.

Quality evaluation of gene function predictions is not trivial and is approached by different research groups in different ways. Most often datasets are assessed by comparing the set of predicted functions for a given gene to a Gold Standard consisting of annotations that are assumed to be correct. This assumption of correctness can be based on any number of criteria. Here we used as our Gold Standard dataset all annotations present in Gramene63 that had a non-IEA (non-Inferred by Electronic Annotation) evidence code, i.e. we used only annotations that had some manual curation. This enabled us to assess the 10 genomes shown in Table II. It is perhaps noteworthy that the IEA and non-IEA annotation sets from Gramene63 frequently contain overlaps, indicating that some of the predicted annotations were manually confirmed afterwards by a curator and that in such cases, a new annotation was asserted with the new evidence code rather than simply upgrading the evidence code from IEA to some other code, thus preserving the IEA annotations in Gramene63 that are produced by the Ensembl analysis pipeline [56], a requirement for comparing GOMAP-produced IEA datasets to the IEA datasets produced by the Ensembl pipeline.

A general limitation of using Gold Standards for quality evaluation is that they can never be assumed to be complete, and therefore false positives in the prediction cannot be distinguished from false negatives in the Gold Standard. In other words, is gene X, function Y truly a wrong prediction or has it simply not yet been discovered experimentally? This problem is laid out in more detail in [57]. As a consequence, the quality of larger prediction sets will be systematically underestimated compared to smaller ones, and this effect is exacerbated the more incomplete the Gold Standard is.

There are many different metrics that have been used to evaluate the quality of predicted functional annotations. For the maize B73 GOMAP annotation assessment in [12], we had used a modified version of the hierarchical evaluation metrics originally introduced in [58] because they were simple, clear, and part of an earlier attempt at unifying and standardizing GO annotation comparisons [59]. In the meantime, Plyusnin et al. published an approach for evaluating different metrics showing variation among the robustness of different approaches to quality assessment [60]. Based on their recommendations, we use here the SimGIC2 and Term-centric Area Under Precision-Recall Curve (TC-AUCPCR) metrics. We also evaluated with the F_max_ metric, simply because it is widely-used (e.g., by [10]), even though according to Plyusnin et al., it is actually a flawed metric [60]. Results of the quality assessments for the 10 genomes where a Gold Standard was available are shown in Table III and Figure S2. While evaluation values differ between metrics and the scores are not directly comparable, a few consistent patterns emerge: GOMAP annotations are almost always better than Gramene and Phytozome annotations in the Cellular Component and Molecular Function aspect, with the only three exceptions being the Molecular

**Table III:** Qualitative metrics of functional annotation sets predicted by GOMAP, Gramene, and Phytozome. This table is visualized in Figure S2.

| Genome | Dataset | SimGIC2 |  |  | TC-AUCPCR |  |  | Fmax |  |  |
| --- | --- | --- | --- | --- | --- | --- | --- | --- | --- | --- |
|  |  | CC | MF | BP | CC | MF | BP | CC | MF | BP |
| *Brachypodium distachyon* | GOMAP | 0.404149 | 0.464127 | 0.223830 | 0.233442 | 0.230701 | 0.118526 | 0.741361 | 0.740897 | 0.526881 |
|  | Gramene63 (IEA only) | 0.317801 | 0.420859 | 0.349406 | 0.129163 | 0.192507 | 0.111361 | 0.691016 | 0.738542 | 0.650325 |
|  | Phytozome12 | 0.370264 | 0.370521 | 0.352206 | 0.112582 | 0.136832 | 0.085628 | 0.717759 | 0.697076 | 0.660603 |
| *Hordeum vulgare* | GOMAP | 0.400087 | 0.470012 | 0.238177 | 0.237231 | 0.261399 | 0.130784 | 0.745272 | 0.750213 | 0.560096 |
|  | Gramene63 (IEA only) | 0.306119 | 0.426601 | 0.381010 | 0.157352 | 0.228797 | 0.136002 | 0.680996 | 0.742638 | 0.665696 |
| *Medicago truncatula* A17 | GOMAP | 0.371795 | 0.451258 | 0.213407 | 0.272809 | 0.282650 | 0.139032 | 0.730838 | 0.726991 | 0.531406 |
|  | Gramene63 (IEA only) | 0.329600 | 0.437274 | 0.343561 | 0.176497 | 0.265887 | 0.133503 | 0.701093 | 0.749900 | 0.654297 |
|  | Phytozome12 | 0.358311 | 0.367257 | 0.363013 | 0.144247 | 0.170863 | 0.110386 | 0.717307 | 0.698429 | 0.661233 |
| *Oryza sativa* | GOMAP | 0.408945 | 0.482650 | 0.248207 | 0.298502 | 0.303384 | 0.159724 | 0.751121 | 0.757181 | 0.559221 |
|  | Gramene63 (IEA only) | 0.328761 | 0.423191 | 0.341193 | 0.167619 | 0.265410 | 0.135451 | 0.711309 | 0.738732 | 0.643827 |
|  | Phytozome12 | 0.049975 | 0.041007 | 0.044279 | 0.000003 | 0.000003 | 0.000002 | 0.470134 | 0.266628 | 0.239256 |
| *Sorghum bicolor* | GOMAP | 0.404852 | 0.466708 | 0.224011 | 0.316873 | 0.337380 | 0.169883 | 0.746540 | 0.742001 | 0.534258 |
|  | Gramene63 (IEA only) | 0.323037 | 0.400241 | 0.353135 | 0.177038 | 0.260198 | 0.154157 | 0.711107 | 0.712170 | 0.653591 |
|  | Phytozome12 | 0.356091 | 0.348264 | 0.340124 | 0.151947 | 0.177579 | 0.110483 | 0.715714 | 0.675147 | 0.641535 |
| *Triticum aestivum* | GOMAP | 0.410582 | 0.489881 | 0.229271 | 0.050762 | 0.030610 | 0.019360 | 0.736476 | 0.762420 | 0.533897 |
|  | Gramene63 (IEA only) | 0.362452 | 0.476685 | 0.395112 | 0.040992 | 0.043701 | 0.027872 | 0.737769 | 0.762059 | 0.670953 |
| *Zea mays* B73.v4 | GOMAP | 0.417455 | 0.467339 | 0.245373 | 0.302761 | 0.290371 | 0.153011 | 0.759504 | 0.746870 | 0.564707 |
|  | Gramene63 (IEA only) | 0.303231 | 0.416301 | 0.346308 | 0.175735 | 0.250075 | 0.138275 | 0.662987 | 0.732860 | 0.647725 |
| *Zea mays* Mo17 | GOMAP | 0.399521 | 0.464265 | 0.225632 | 0.236209 | 0.239598 | 0.125599 | 0.744360 | 0.743026 | 0.537489 |
| *Zea mays* PH207 | GOMAP | 0.394481 | 0.436266 | 0.224226 | 0.221709 | 0.221266 | 0.117086 | 0.743111 | 0.718933 | 0.533092 |
| *Zea mays* W22 | GOMAP | 0.397602 | 0.463499 | 0.223511 | 0.210198 | 0.217609 | 0.113262 | 0.743783 | 0.742341 | 0.535572 |

Download this table \(CSV\)

Function aspect for *T. aestivum* using the TC-AUCPCR and the F_max_ metric and the Cellular Component aspect for *M. truncatula* A17 using the F_max_ metric. Conversely, GOMAP predictions achieve consistently lower quality scores in the Biological Process aspect with the exception of *B. dystachion*, *O. sativa*, and *S. bicolor* with the TC-AUCPR metric. Generally, annotations that are better in one aspect are also better in the other two aspects but the ranking of annotations does not necessarily hold across metrics. The Phytozome annotation for *O. sativa* is an outlier in terms of its comparative quality, potentially because it is based on a modified structural annotation that differs substantially from the Gold Standard and the other annotations under comparison.

### C. Phylogenetic Tree Analyses

With the comparative quality of gene function predictions in hand, we approached the question of whether the datasets could be used together for comparative functional analysis across all genomes. As a simple first step, we began to work toward understanding the degree to which trees built based on gene functions agree with known, well-documented evolutionary relatedness. We constructed neighbor-joining and parsimony trees of the 18 plant genomes, and visulized them using iTOL [61]. The two tree topologies, rooted at *P. lambertiana*, were compared to one another and to the topology of the expected tree (Figure 2). For both the neighbor-joining (Figure 3a) and parsimony trees (Figure 3b), one common difference is noted: *S. bicolor* is not at the base of the *Z. mays* clade as expected, and is clustered with *B. distachyon* instead. Notable differences between the neighbor-joining and parsimony tree are the following: *C. sativa* appears at the base of the eudicots instead of *G. raimondii* in the neighbor-joining tree, while *G. raimondii* is grouped with *C. sativa* and *A. hypogaea* is grouped with *G. max* in the parsimony tree. Second, *O. sativa* was expected to be at the base of the BOP clade, but appears at the base of *Z. mays* in the neighbor-joining tree, and at the base of all angiosperms in the parsimony tree. Differences among relationships within the *Z. mays* clade constaining B73, PH207, W22, and Mo17 were disregarded given the high degree of similarity across annotation sets and the fact that these relationships are not clear given the complex nature of within-species relationships.

**Figure 3:**
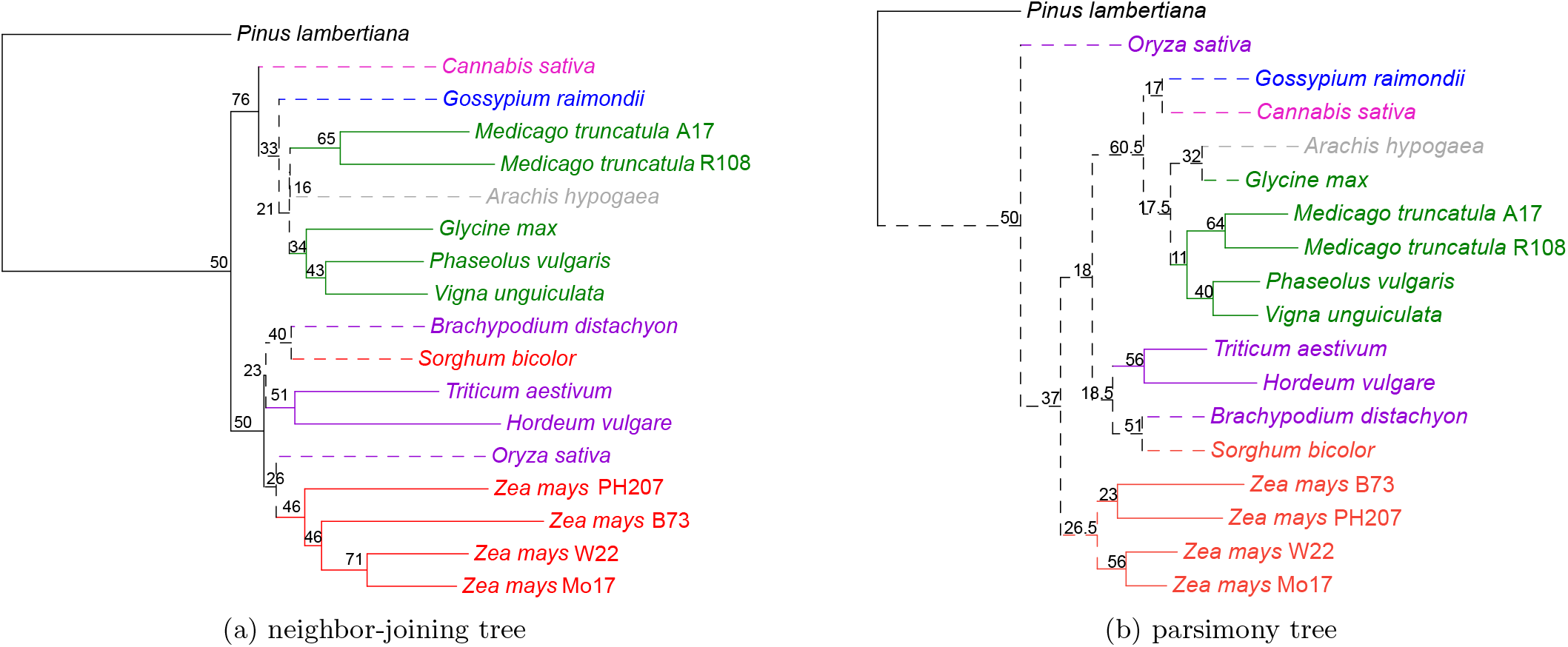
Neighbor-joining and parsimony trees. Phylograms are colored and rooted as described in Figure 2. For both neighbor-joining (a) and parsimony (b), node values represent the jackknifing support values derived by removing 40% of GO terms in the dataset. Dashed lines mark deviations from known phylogenetic relationships. Tree scales are shown above each, with NJ showing distances and parsimony showing changes in character state.

Due to differences between the function-based dendrograms and the expected tree, jackknifing analysis was carried out by removing terms (columns in underlying datasets) to determine the degree to which the underlying datasets support specific groupings based on functional term assignments. This analysis was carried out for both neighbor-joining and parsimony trees. First, trees were generated by omitting 5% to 95% of the dataset in increments of 5 to determine the threshold at which the tree topologies deviated from those generated using the full dataset. That threshold was reached at 45% for both neighbor-joining and parsimony; therefore, we used trees generated with 40% of the data removed for reporting branch support values for the topology (Figure 3). Comparing the two trees, the parsimony topology was not as solid as that of the neighbor-joining at jackknife values up to 40%. Based on this robustness for neighbor-joining treebuilding in general, we carried out all subsequent analyses using neighbor-joining treebuilding methods.

We considered investigating the effect of using one GO aspect to generate our neighbor-joining tree. In other words, we generated the neighbor-joining trees using cellular component GO terms, molecular function GO terms, and biological process GO terms separately (Supplementary Figure 3). Of the 14,303 total GO terms, 1,524 are cellular component terms, 3,926 are molecular function terms, and 8,853 are biological process terms. Out of the three single aspect phylogenetic trees, the one built using molecular function terms is the closest to our neighbor-joining tree obtained using all GO terms in our datasets Figure 3a. The only difference is that *A*. *hypogaea* and *G. max* are clustered in the molecular function tree, while they are not in our neighbor-joining tree Figure 3a. In the cellular component tree, *G. raimondii* and *C. sativa* are clustered together when they are not in the neighbor-joining tree with all GO aspects Figure 3a. Also, *O. sativa* is at the base of the monocots just like in the expected tree, but not in the neighbor-joining tree Figure 3a. In the biological process tree, *O. sativa* is at the base of the angiosperms and there is no clear separation between monocots and dicots. In all the three single aspect phylogenetic trees and our all-aspect neighbor-joining tree, *A. hypogaea* is never placed at the base of the NPAAA clade. Also, *B. distachyon* and *S. bicolor* are always clustered together. Overall, the topologies constructed using one GO term aspect at a time are close to that of our neighbor-joining tree, such that not one GO term aspect alone restored the topology of the expected tree.

To map the source of discrepancies to specific gene annotation sets, we generated various neighbor-joining trees excluding one genome each time, an additional tree with both *Medicago* genomes excluded simultaneously, and another with all *Z. mays* genomes excluded simultaneously. To exemplify this, see the monocot clade in Figure 2 and the lower (monocot) clade in Figure 3a. When the neighbor-joining tree was generated, two species are misplaced: *S. bicolor* and *O. sativa*. As shown in Figure 5a, removal of *O. sativa* corrects one error (itself) but does not correct the errant grouping of *S. bicolor* with *S. distachyon*. In Figure 5b, it is shown that the removal of *S. bicolor* corrects the errant grouping of itself and *B. distachyon*, but *O. sativa* placement remains incorrect. However, as shown in Figure 5c, the removal of *B. distachyon* generates a tree where all relationships are in agreement with known species-level relationships. (Note well: all individual annotation sets were progressively removed, not just the three shown in the example.)

With this observation in hand, we sought to determine the minimum number of genomes that could be removed to create a tree that matched the expected tree topology. All possible combinations of removing 0-5 genomes to restore the topology were tested, and 10 combinations of minimum amount of genomes to be removed were obtained. The removal of four genomes was required to generate function-based trees consistent with known phylogenetic relationships. Of the 10, we selected the one that had the genomes that were most frequently part of a solution (*O. sativa*, 8; *B. distachyon*, 7; *C. sativa*, 6; *A. hypogaea*, 5; *S. bicolor*, 4; *G. raimondii*, 4; *G. max*, 4; *T. aestivum*) to show in this paper (the other combinations can be found in our publicly available dataset). To elaborate, the genomes removed here are *O. sativa*, *B. distachyon*, *C. sativa*, and *A. hypogaea* (Figure 4). Jackknifing analysis was also carried out for this dataset with support shown. Branch support is generally higher than that for the full dataset (i.e., branch support is higher in Figure 4 than in Figure 3a), and removing genomes that are causing variations seem to stabilize the tree.

**Figure 4:**
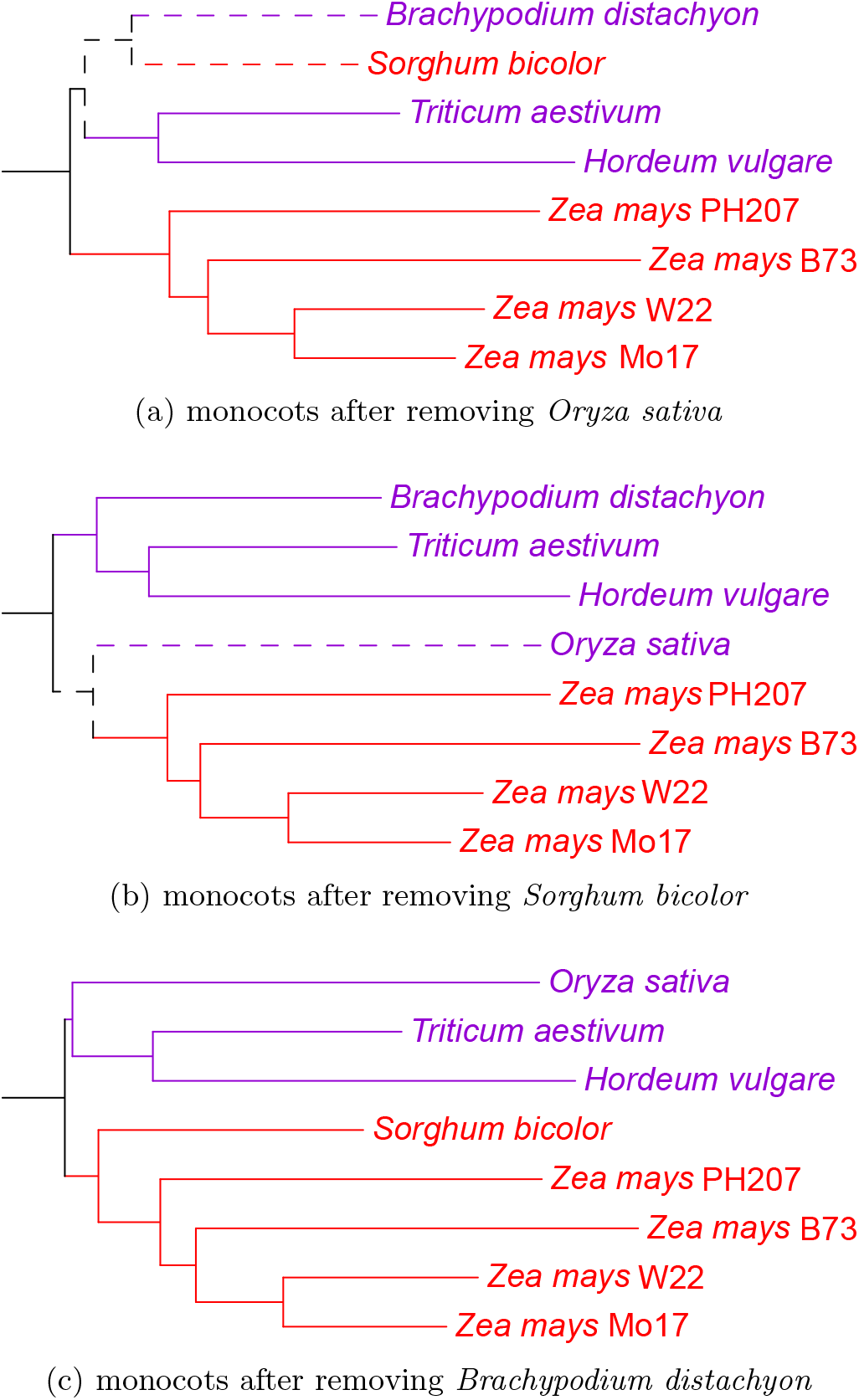
Restoring monocot relationships. Phylograms are colored and rooted as described in Figure 2. Dashed lines mark deviations from known phylogenetic relationships. Monocot topology changes with removal of a single species: (a) *O. sativa*, (b) *S. bicolor*, and (c) *B. distachyon*. Tree scale is shown above

**Figure 5:**
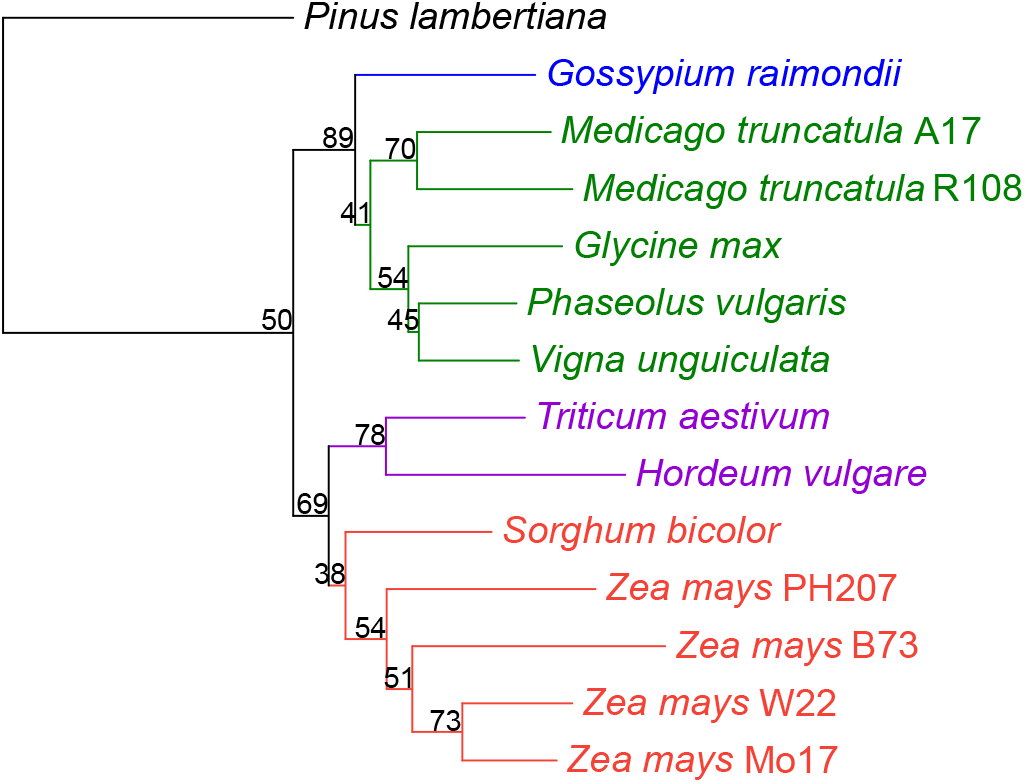
Restoring known phylogentic relationships to the NJ tree via removal of a minimal number of species. Phylograms are colored and rooted as described in Figure 2. Node values represent the jackknifing support values derived by removing 40% of GO terms in the dataset. 4 genomes have been removed: *C. sativa*, *O. sativa*, *B. distachyon*, and *A. hypogaea*. Tree scale is shown above.

### D. Potential Causes of Unexpected Groupings

As a first step toward explaining discrepancies between known evolutionary relationships and those resulting from comparative analysis of genome-wide gene function predictions, we assessed the quality of each genome assembly and structural annotation set using GenomeQC [62]. Tables IV–V and Figures 6–7 represent the resulting assembly quality, structural annotation measures of quality, and proportion of single-copy BUSCOs (Bench-marking Universal Single-Copy genes) [63] that were generated. Although these analyses make evident that the species annotated are comparatively different in both natural genome characteristics and in assembly and annotation quality aspects, it is not the case that the four species responsible for deviations between the functional annotation dendrograms and known phylogenetic relationships (i.e., *C. sativa*, *A. hypogaea*, *O. sativa* and *B. distachyon*) create these discrepancies due to issues of genome assembly and/or annotation quality. One potential for some explanation is in relation to *C. sativa*, which is the only genome that has an assembly length larger than the expected (see IV), and a comparatively large proportion of missing BUSCOs in the assembly (see Figure 6). Similarly, for *A. hypogaea* and *O. sativa*, there is a large proportion of missing BUSCOs in the annotations (see Figure 7).

**Table IV:**
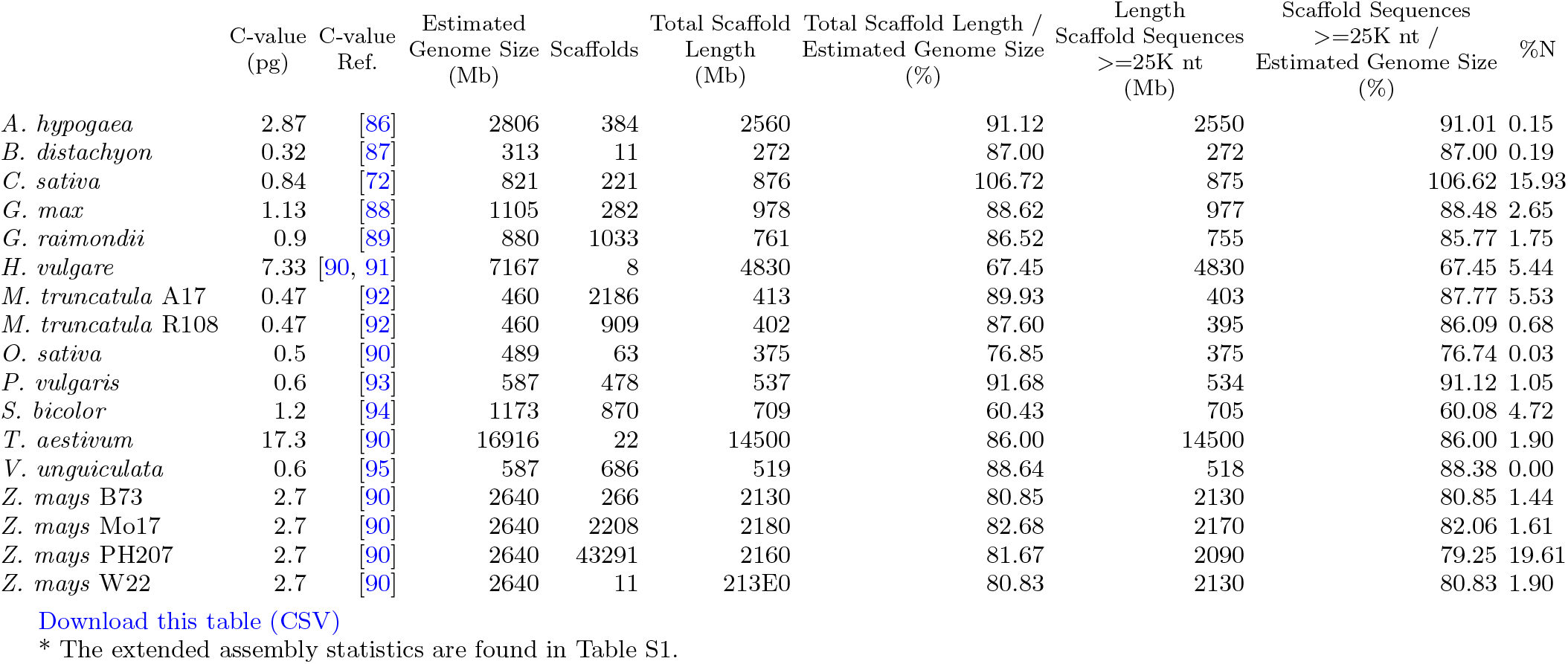
Assembly statistics*

**Table V:** Structural annotation table directly from GenomeQC.

|  | Gene Models | Min.<br>Gene Length<br>(bp) | Max.<br>Gene Length<br>(bp) | Avg.<br>Gene Length<br>(bp) | Gene Models<br><200 bp | Transcripts | Avg. Transcripts<br>per Gene Model | Exons | Avg. Exons<br>per Transcript | Avg. Exon<br>Length (bp) |
| --- | --- | --- | --- | --- | --- | --- | --- | --- | --- | --- |
| *A. hypogaea* | 67128 | 163 | 342359 | 3972.3 | 48 | 84714 | 1.3 | 423763 | 4.8 | 296.4 |
| *B. distachyon* | 36647 | 90 | 47411 | 3012.5 | 175 | 47917 | 1.3 | 263865 | 5.5 | 314.1 |
| *C. sativa* | 29815 | 21 | 976063 | 3450.0 | 2187 | 34876 | 1.2 | 234157 | 6.6 | 291.9 |
| *G. max* | 52872 | 90 | 94733 | 4017.0 | 1363 | 86256 | 1.6 | - | - | - |
| *G. raimondii* | 37505 | 90 | 51175 | 3243.5 | 71 | 77267 | 2.1 | 527563 | 6.7 | 271.3 |
| *H. vulgare* | - | - | - | - | - | 248270 | - | 1715898 | - | 278.9 |
| *M. truncatula* A17 | 51541 | 60 | 102191 | 2566.9 | 1985 | 74213 | 1.4 | 318421 | 4.4 | 313.2 |
| *M. truncatula* R108 | 55706 | 72 | 62996 | 2232.4 | 2404 | 61019 | 1.1 | 220904 | 3.6 | 270.6 |
| *O. sativa* | 35825 | 72 | 304271 | 3098.7 | 57 | 42378 | 1.2 | 192499 | 4.5 | 350.3 |
| *P. vulgaris* | 27433 | 93 | 90772 | 3943.8 | 108 | 36995 | 1.3 | - | - | - |
| *S. bicolor* | 34129 | 96 | 88337 | 3713.2 | 56 | 47121 | 1.4 | 266301 | 5.6 | 350.9 |
| *T. aestivum* | 107891 | 54 | 124945 | 3488.9 | 686 | 133744 | 1.2 | 749233 | 5.8 | 303.3 |
| *V. unguiculata* | 29773 | 90 | 81066 | 3880.9 | 408 | 42287 | 1.4 | - | - | - |
| *Z. mays* B73 | 39324 | 111 | 128402 | 4117.8 | 38 | 151959 | 3.9 | 1488381 | 9.7 | 281.5 |
| *Z. mays* Mo17 | 38620 | 21 | 146217 | 4076.8 | 1225 | 46530 | 1.2 | 272323 | 5.9 | 297.1 |
| *Z. mays* PH207 | 40557 | 111 | 714207 | 5993.3 | 53 | 40557 | 1.0 | 208149 | 5.1 | 252.1 |
| *Z. mays* W22 | 40691 | 87 | 154495 | 4330.4 | 64 | 51717 | 1.3 | 313830 | 6.1 | 292.4 |

Download this table \(CSV\)
Dashes (-) represent missing data.
Missing data in *H. vulgare* is due to the absence of gene model information in the GFF file.
Missing data in *G. max*, *P. vulgaris*, and *V. unguiculata* is due to the absence of exon information in their GFF files.

**Figure 6:**
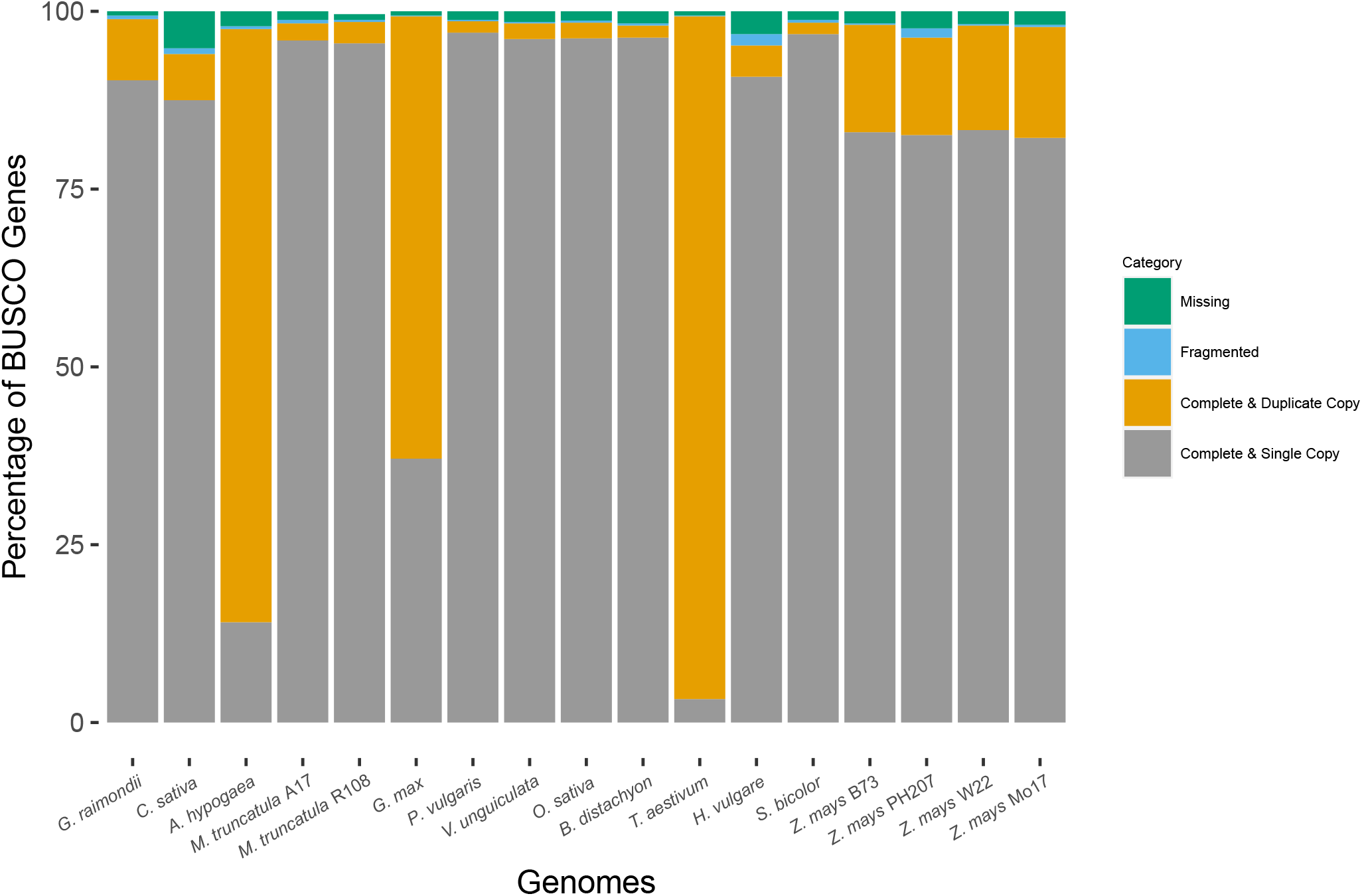
Assembly BUSCO plot generated using GenomeQC. Genomes analyzed are shown across the X-axis, and are ordered to match the occurrence of species shown in Figure 2. Percentage of BUSCO genes across four gene categories are stacked, with each adding up to 100 percent (Y-axis). Complete and single-copy genes are shown in grey, complete and duplicated copies in orange, fragmented copies in blue, and missing are shown copies in green.

**Figure 7:**
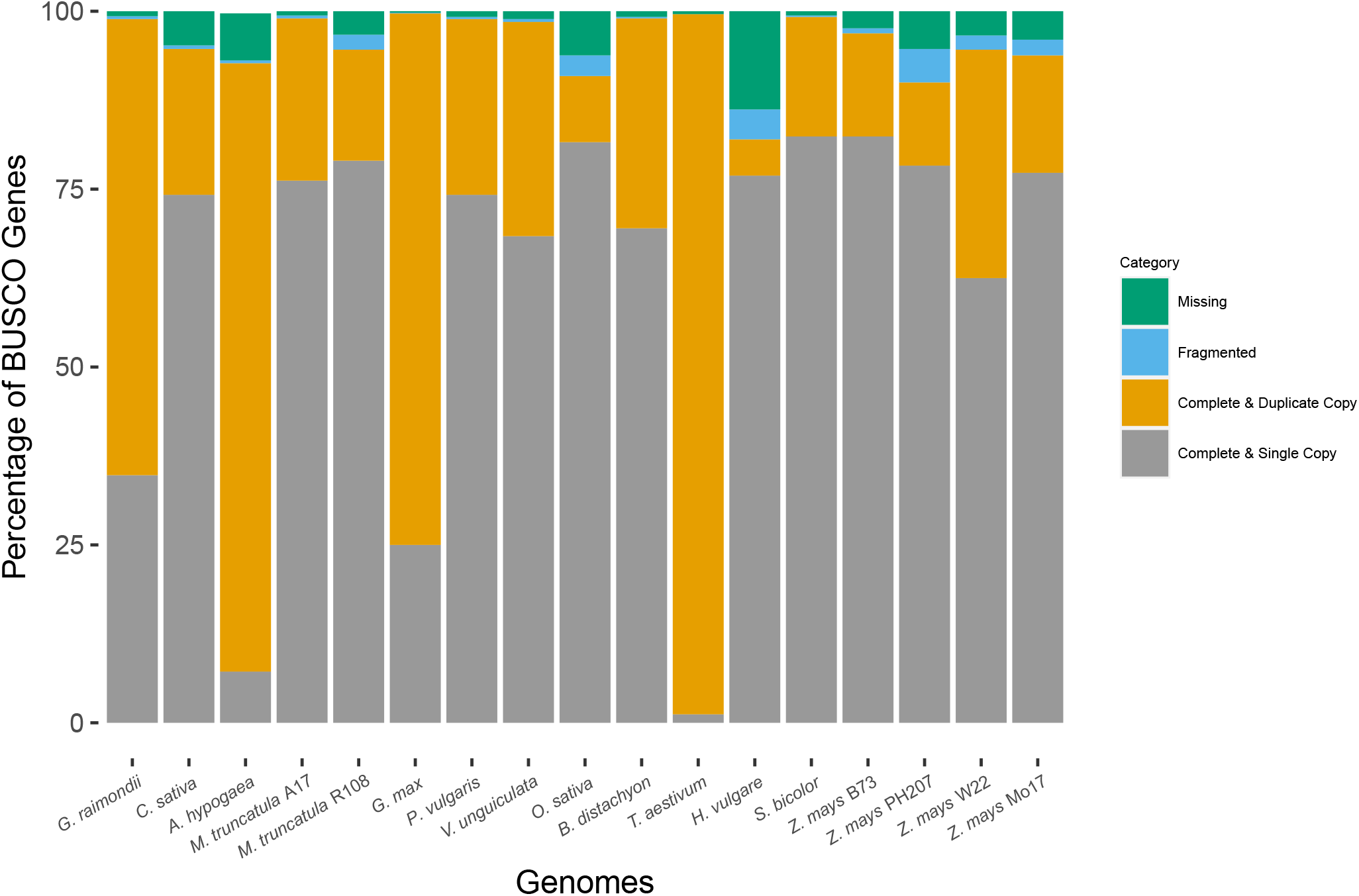
Annotation BUSCO plot generated using GenomeQC. Genomes analyzed are shown across the X-axis, and are ordered to match the occurrence of species shown in Figure 2. Percentage of BUSCO genes across four gene categories are stacked, with each adding up to 100 percent (Y-axis). Complete and single-copy genes are shown in grey, complete and duplicated copies in orange, fragmented copies in blue, and missing are shown copies in green.

## III. DISCUSSION

In this study, we used the GOMAP pipeline to produce whole-genome GO annotations for 18 genome assembly and annotation sets from 14 plant species [9]. Assessments of the number of terms predicted as well as the quality of predictions indicate that GOMAP functional prediction datasets cover more genes, contain more predictions per gene, and are of similar quality to prediction datasets produced by other systems, thus supporting the notion that these high-coverage datasets are a useful addition for researchers who are interested in genome-level analyses, including efforts aimed at prioritizing candidate genes for downstream analyses. Given that we can now produce high-quality, whole genome functional annotations for plants in a straightforward way, we intend to produce more of these over time (indeed we recently annotated *Vitis vinifera* [64], *Brassica rapa* [65], *Musa acuminata*[66], *Theobroma cacao* [67], *Coffea canephora* [68], *Vaccinium corymbosum* [69] *Solanum lycopersicum [70]*, and *Solanum pennellii* [71]).

With 18 genome functional annotations in hand, we sought to determine whether and how researchers could use multispecies GO annotation datasets to perform comparative functional genomics analyses. As a proof of concept, we adapted phylogenetic tree-building methods to use the gene function terms assigned to genes represented by the genomes to build dendrograms of functional relatedness and hypothesized that if the functions were comparable across species, the resulting trees would closely match evolutionary relationships. To our delight and surprise, the neighbor-joining and parsimony trees (Figure 3) did resemble known phylogenies, but were not exact matches to broadly accepted phylogenetic relationships. After removing the minimum number of genomes that resulted in restoration of the expected evolutionary relationships, we found that the individual species that may be responsible for the discrepancies observed in Figure 3 were *C. sativa*, *A. hypogaea*, *O. sativa* and *B. distachyon*. We hypothesize that the following could account for such errant relationships:

1. Quality of sequencing and coverage assembly: genomes of similarly high sequence coverage that have excellent gene calling would be anticipated to create the best source for functional annotation. Genomes of comparatively lower, or different, character would be anticipated to mislead treebuilding and other comparative genomics approaches.
2. Shared selected or natural traits: species that have been selected for, e.g., oilseeds may share genes involved in synthesis of various oils. Other shared traits would be anticipated to cause similarities for species with those shared traits.
3. Lack of good representation of diverse plant biology aspects in the GO graph: most plant-specific GO terms were derived from functional analysis of one model species, *Arabidopsis thaliana*. This single-source for presence of plant-specific functions limits the graph from containing unique functional aspects of plant biology represented in other species’ genomes.
4. Use of a simple method of treebuilding based on the presence or absence of gene function terms: the method we devised and describe here is not sophisticated enough to make full use of information in the GO graphs such that we recover the full detail of the species’ evolutionary histories from the simple method.

To consider the first of these, we looked at genome assembly and annotation quality metrics (see Tables IV–V and Figures 6–7). For *B. distachyon* we could find no compelling evidence that assembly structural or functional annotation quality differed significantly from all others, except in the case of *C. sativa*, where we noted that the assembly length exceeded the predicted genome size based on C-values for genome sizes reported previously [72]. In this case, the fact that the *C. sativa* line sequenced is not inbred [73] may be responsible for the inflated assembly size relative to what is expected. This means that in the assembly, there are likely regions where alleles between chromosomes do not align, which would inflate the overall length of the assembly. In addition, the assembly misses a large proportion of BUSCO genes compared to most other genomes included in this analysis. Indeed, the comparatively low-quality assembly for *Cannabis* genome has been noted by others [74], and our preliminary investigations indicate that the assembly length is in fact longer than expected.

In an attempt to better understand conflicting phylogenetic signals that could be caused by the second potential cause, i.e., shared selected or natural traits, we mapped all GO terms that exist in our binary matrix and traced character history (presence/absence) on the nodes and leaves of our expected evolutionary tree using the software Mesquite version 3.61 [75]. These data summarize the gain or loss of each GO term across the species described in this paper and can be found in our Github repository. We carried out a number of simple experiments to reveal which terms could be causal for errant relationships (e.g., dropping all unique terms from the *B. distachyon* dataset, reconstructing the term states at nodes that should be where *B. distachyon* should occur, etc., and could not identify any biologically compelling patterns. (Because these analyses were not fruitful, they were not specifically included in our materials and methods, though we do include the input datasets here, at the link provided in section V “Availability of Source Code and Supporting Data” of the paper, for others to consider and peruse independently.)

An important limitation that must be mentioned is the effect of the third potential cause of errant on our generated phylogenetic trees: a deficiency of terms describing plant biology. Because most GO terms specific to plant biology are likely derived from *Arabidopsis*, a model dicot species, gene functions unique to other species are expected to be missing from the GO graphs [76]. This source of error will only be corrected over time as gene functions unique to diverse plant species are populated into the GO graph.

We consider the most likely explanation for observed discrepancies between the known evolutionary phylogenies and dendrograms created based on GO terms describing gene function to be a result of the fourth explanation: the simplicity of the treebuilding models and methods we used for these analyses. Because the tree-building and analytics described in this paper were based on the presence/absence of GO terms, novel terms are highly influential to the outcomes of the analysis and the number of times a term is used does not influence the outcome at all. In contrast, plant genomes are notable for having many duplicated genes as a result of whole-genome and segmental duplications over evolutionary history, so these duplications are in fact a feature of and marker for what happened to that genome over time. Therefore, using presence/absence of GO terms where the count of term occurrences are not weighted may be too simple to get at the genuine biological complexities represented in any given plant genome. Our simplistic demonstration of the utility of GO datasets for comparative functional genomics shows that more sophisticated methods are very promising for comparative functional genomics analyses.

It should be noted that comparative analyses using gene functions are not completely absent from the literature - though they are absent for large genome comparisons in plants. An example of such existing comparative use of GO is one where a given tree topology was used to look for gains and losses of functions mapped to independently derived trees, which was reported by Schwacke et al. [77]. They report, as an example of their method, an analysis of gene loss in *Cuscuta*, a parasitic plant based on analysis using the Mapman ontology. In their work, they showed considerable loss of genes, which is a hallmark of the parasitic lifestyle. Our efforts differ in that we used the functions directly to infer tree structures as a demonstration that sufficient biological signal is present in GO-based datasets of genome-wide function prediction to reproduce known biological relationships. The method we used was quick and dirty, and we anticipate that refinements in approach that consider multiple copies of genes, as well as using different types of graph and network representations beyond tree structures, are logical next steps for refining the use of GO terms for comparative functional genomics analyses in plants. With that in mind, we look forward not only to developing systems to support GO-based comparative functional genomics tools, but also to seeing the tools other research groups will develop to approach the use of these datasets to formulate novel comparative functional genomics hypotheses.

## IV. METHODS

### A. Acquiring Input Datasets

For each of the 18 genomes listed in Table I, information on how to access input annotation products are listed by DOI. For each, one representative translated peptide sequence per protein coding gene was selected and used as the input for GOMAP, a gene function prediction tool for plants that is actively maintained, updated, and versioned. Details of how GOMAP annotations are derived including the specificity of component datasets and which terms are retained are described elsewhere [12] [9]. In brief, GOMAP annotations are a combination of the annotations from multiple sources. GOMAP combines the annotations from all the sources and removes the less specific annotations that could be inferred from the more specific annotations, keeping only the most specific terms for each gene that cannot be inferred from other terms (i.e., only leaf terms). Unless the authors of the genome provided a set of representative sequences designated as canonical, we chose the longest translated peptide sequence as the representative for each gene model. In general, non-IUPAC characters and trailing asterisks (*) were removed from the sequences, and headers were simplified to contain only non-special characters. The corresponding script for each dataset can be found at the respective DOI. Based on this input, GOMAP yielded a functional annotation set spanning all protein-coding genes in the genome. Using the Gene Ontology version releases/2020-10-09, this functional annotation set was cleaned up by removing duplicates, annotations with qualifiers (NOT, contributes to, colocalizes with; column 4 in the GAF 2.1 format), and obsolete GO terms. Any terms containing alternative identifiers were merged to their respective main identifier, uncovering a few additional duplicates, which were also removed. Table SII shows the number of annotations removed from each dataset produced.

To compare the quality of GOMAP predictions to currently available functional predictions from Gramene and Phytozome, we downloaded IEA annotations from Gramene (version 63, [50], https://www.gramene.org/) and Phytozome (version 12, [51], https://phytozome.jgi.doe.gov/) for each species with functional annotations of the same genome version. These datasets were cleaned as above. Similarly cleaned non-IEA annotations from Gramene63 served as the Gold Standard wherever they were available. More detailed information on how these datasets were accessed can be found at https://github.com/Dill-PICL/GOMAP-Paper-2019.1/blob/master/data/go_annotation_sets/README.md.

### B. Quantitative and Qualitative Evaluation

The number of annotations in each clean dataset was determined and related to the number of protein coding genes (based on transcripts in the input FASTA file). This was done for separately for each GO aspect as well as in total.

The ADS software version published in [60] is available from https://bitbucket.org/plyusnin/ads/. We used version b6309cb (also included in our code as a submodule) to calculate SimGIC2, TC-AUCPCR, and F_max_ quality scores. To provide the information content required for the SimGIC2 metric, the Arabidopsis GOA from https://www.ebi.ac.uk/GOA/arabidopsis_release was used in version 2021-02-16.

### C. Cladogram Construction

For clustering, we first collected all GO terms annotated to any gene in each genome into a list and removed the duplicates, yielding a one-dimensional set of GO terms for each genome (*T*). Next, we added all parental terms for each term in this set (connected via *is a* in the ontology), their respective parental terms and higher, recursively continuing up to the very root of the ontology. Then we once again removed the duplicates, yielding a set *S* containing the original terms from set *T* as well as all terms proximal to them in the Gene Ontology directed acyclical graph. These sets with added ancestors served as a starting point of our tree-building analyses: pairwise distances between the genomes were calculated using the Jaccard distance as a metric of the dissimilarity between any two sets *a* and *b*.

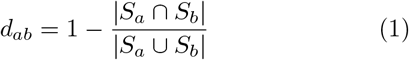

A neighbor-joining tree was constructed based on the generated pairwise distance matrix using PHYLIP [52]. Additionally, term sets *S* of all genomes were combined into a binary matrix (with rows corresponding to genomes and columns corresponding to GO terms, values of 0 or 1 indicating whether a term is present or absent in the given set). PHYLIP pars was used to construct a parsimony tree from this binary matrix.

*P. lambertiana*, a gymnosperm, was included in the dataset as an outgroup to the angiosperms to separate between the monocot and eudicot clades. iTOL [78] was used to visualize the trees using their Newick format, and root them at *P. lambertiana*. Moreover, a cladogram representing the known phylogeny of the included taxa was created by hand based on known evolutionary relationships [79, 80, 81, 82, 83]. This was used to compare the generated phylogenetic relationship based on functional similarity with the evolutionary relationships of the plant genomes.

Jackknifing analysis was carried out for both parsimony and neighbor-joining trees to assess the support for each clade based on the proportion of jackknife trees showing the same clade. To this end, 40% of the terms in *T* were randomly removed, ancestors of the remaining terms were added and trees constructed as above. The majority rule consensus tree of 100 individual trees was calculated with the jackknife values represented on each branch. The tree was then visualized using iTOL using its Newick format, and rooted again at *P. lambertiana*.

### D. Genome Quality Evaluation

Genome size was estimated from the C-values obtained from the Plant DNA C-values data resource from the Kew Database (https://cvalues.science.kew.org). The mean C-value for a given species was used for calculating genome size estimates in base pairs (bp) using the method of [84]. In brief,

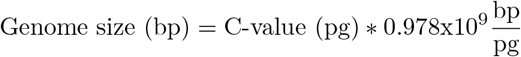

The estimated genome size (listed in Table IV) was used as an input for GenomeQC (https://genomeqc.maizegdb.org/) [62] to calculate quality metrics. For genomes that were too large to submit through the GenomeQC webtool or had missing exon information, modified scripts of those found in GitHub of GenomeQC (https://github.com/HuffordLab/GenomeQC, commit e6140ee) were applied to calculate the assembly and structural annotation metrics in Table IV and 5. BUSCO version 5.0.0 [85] was used to calculate the assembly and annotation BUSCO scores, shown in Figures 6 and 7. The input for assembly BUSCO scores were chromosome sequences, whereas inputs were transcript/mRNA/CDS sequences for the annotation BUSCO scores. For the lineage parameter, the lineage datasets used were as follows: Eudicots for *C. sativa* and *G. raimondii*, Fabales for *A. hypogaea*, *M. truncatula* A17 and R108, *P. vulgaris*, *G. max*, and *V. unguiculata*, and Poales for *B. distachyon*, *O. sativa*, *T. aestivum*, *H. vulgare*, *S. bicolor* and *Z. mays* B73, Mo17, W22 and PH207.

## V. AVAILABILITY OF SOURCE CODE AND SUPPORTING DATA

All data and source code generated are freely available at https://github.com/Dill-PICL/GOMAP-Paper-2019.1 under the terms of the MIT license. All software requirements and dependencies are packaged into a Singularity container so no other setup is required to reproduce our results. We will provide a DOI through Zenodo for the final version of the manuscript after reviews and corrections are incorporated.

An up-to-date list of all available annotation sets can be found at https://dill-picl.org/projects/gomap/gomap-datasets/.

## VI. DECLARATIONS

### C. Author’s Contributions

LF, DP, CFY, KOC, HAD, PJ, DCS, HV, and KW generated annotations for plants as described in this paper. DP and CJLD co-conceived the idea for phylogenetic analysis. DP wrote the code for the analyses in this paper. LF worked with DP to create dendrograms and compare those to phylogenetic trees. LF carried out assembly and annotation metric comparisons. LF, DP, and CJLD wrote the manuscript. All authors read, offered suggestions to improve, and approved the final copy of the manuscript.

## Supporting information

Full Supplement, includes Supplemental Tables

## VII. ACKNOWLEDGEMENTS

Thanks to Steven Cannon for help to understand phylogenetic relationships among eudicots and helpful discussions and to Toby Kellogg and Jedrzej Szymanski for discussions and ideas on how to consider the data. Thanks to Nancy Manchanda for reviewing documentation and for checking genome versions used as input for GenomeQC. Thanks to Darwin Campbell who guided the deposition of datasets with CyVerse for release and DOI assignment. Thanks to the reviewers for their suggestions which have improved this manuscript.

## VIII. AUTHORS’ INFORMATION

KW created the GOMAP system during his time as a graduate student at Iowa State University. LF, DP, CFY, KOC, HV, and PJ are currently graduate students. HAD and DCS are undergraduate students. Each graduate and undergraduate student annotated at least one genome over the course of a research rotation lasting no more than one semester. CJLD coordinated research activities and manuscript preparation.

