## Supplementary material for "Standardized genome-wide function prediction enables comparative functional genomics: a new application area for Gene Ontologies in plants": Full Supplement, includes Supplemental Tables

### IX. SUPPLEMENTARY

Supplementary Table SI: Additional assembly statistics from GenomeQC.

| | Longest Scaffold (bp) | Shortest Scaffold (bp) | Scaffolds >1K nt | Scaffolds >10K nt | Scaffolds >100K nt | Scaffolds >1M nt | Scaffolds >10M nt | N50 | L50 | NG50 | LG50 | %A | %C | %G | %T |
|---|---|---|---|---|---|---|---|---|---|---|---|---|---|---|---|
| *A. hypogaea* | 1.61E+08 | 1000 | 382 | 299 | 42 | 23 | 20 | 1.35E+08 | 9 | 1.35E+08 | 10 | 31.79 | 18.14 | 18.13 | 31.79 |
| *B. distachyon* | 7.55E+07 | 2.88E+04 | 11 | 11 | 5 | 5 | 5 | 5.89E+07 | 3 | 5.89E+07 | 3 | 26.76 | 23.16 | 23.14 | 26.76 |
| *C. sativa* | 1.05E+08 | 6741 | 221 | 218 | 52 | 15 | 10 | 9.19E+07 | 5 | 9.19E+07 | 5 | 27.94 | 14.07 | 14.11 | 27.95 |
| *G. max* | 5.83E+07 | 885 | 281 | 271 | 63 | 20 | 20 | 4.99E+07 | 10 | 4.78E+07 | 11 | 31.77 | 16.91 | 16.91 | 31.77 |
| *G. raimondii* | 7.07E+07 | 1001 | 1033 | 200 | 20 | 14 | 13 | 6.22E+07 | 6 | 6.10E+07 | 7 | 32.81 | 16.32 | 16.32 | 32.81 |
| *H. vulgare* | 7.68E+08 | 2.50E+08 | 8 | 8 | 8 | 8 | 8 | 6.57E+08 | 4 | 5.83E+08 | 6 | 26.27 | 21.02 | 21.01 | 26.26 |
| *M. truncatula* A17 | 5.66E+07 | 1004 | 2186 | 605 | 51 | 8 | 8 | 4.92E+07 | 4 | 4.57E+07 | 5 | 31.60 | 15.66 | 15.65 | 31.57 |
| *M. truncatula* R108 | 3.23E+07 | 1007 | 909 | 481 | 72 | 49 | 18 | 1.28E+07 | 12 | 1.23E+07 | 14 | 33.25 | 16.41 | 16.41 | 33.25 |
| *O. sativa* | 4.33E+07 | 4236 | 63 | 51 | 17 | 12 | 12 | 3.00E+07 | 6 | 2.90E+07 | 8 | 28.21 | 21.77 | 21.78 | 28.21 |
| *P. vulgaris* | 6.30E+07 | 1005 | 478 | 290 | 61 | 13 | 11 | 4.97E+07 | 5 | 4.80E+07 | 6 | 31.84 | 17.61 | 17.64 | 31.86 |
| *S. bicolor* | 8.09E+07 | 1005 | 870 | 232 | 69 | 12 | 10 | 6.87E+07 | 5 | 6.87E+07 | 9 | 26.74 | 20.90 | 20.91 | 26.73 |
| *T. aestivum* | 8.31E+08 | 4.74E+08 | 22 | 22 | 22 | 22 | 22 | 7.10E+08 | 10 | 6.74E+08 | 12 | 26.46 | 22.59 | 22.59 | 26.46 |
| *V. unguiculata* | 6.53E+07 | 2922 | 686 | 679 | 103 | 13 | 11 | 4.17E+07 | 6 | 4.13E+07 | 7 | 33.31 | 16.40 | 16.42 | 33.34 |
| *Z. mays* B73 | 3.07E+08 | 5568 | 266 | 261 | 48 | 12 | 10 | 2.24E+08 | 5 | 1.82E+08 | 6 | 26.18 | 23.09 | 23.10 | 26.19 |
| *Z. mays* Mo17 | 3.06E+08 | 1007 | 2208 | 1864 | 129 | 11 | 10 | 2.20E+08 | 5 | 1.83E+08 | 6 | 26.14 | 23.03 | 23.05 | 26.17 |
| *Z. mays* PH207 | 3.03E+08 | 500 | 16840 | 1115 | 61 | 11 | 10 | 2.15E+08 | 5 | 1.76E+08 | 6 | 21.34 | 18.40 | 18.40 | 21.36 |
| *Z. mays* W22 | 3.11E+08 | 1.25E+07 | 11 | 11 | 11 | 11 | 11 | 2.23E+08 | 5 | 1.83E+08 | 6 | 26.11 | 22.92 | 22.94 | 26.13 |

Download this table (CSV)

N50: The length of the shortest scaffold/contig in the list of L50 sequences.

L50: The number of sequences whose sum of lengths make up 50% or more of the total assembly length.

NG50: The length of the shortest scaffold/contig calculated in the same manner as N50 but based on estimated genome size rather than total assembly length.

LG50: The number of sequences whose sum of lengths make up 50% or more of the estimated genome size.

Supplementary Table SII: Number of removed annotations during cleanup.

| Genome | Dataset | Obsolete Annotations | Duplicates | Annotations with Modifiers |
|---|---|---|---|---|
| *Arachis hypogaea* | GOMAP | 3437 | 13 | 912 |
| *Brachypodium distachyon* | GOMAP | 2512 | 49 | 789 |
| | Gold Standard Gramene 63 (no IEA) | 21 | 204 | 0 |
| | Gramene63 (IEA only) | 166 | 114 | 0 |
| | Phytozome12 | 99 | 18 | 0 |
| *Cannabis sativa* | GOMAP | 1714 | 6 | 757 |
| *Glycine max* | GOMAP | 3333 | 10 | 930 |
| *Gossypium raimondii* | GOMAP | 1781 | 7 | 822 |
| *Hordeum vulgare* | GOMAP | 1877 | 8 | 815 |
| | Gold Standard Gramene 63 (no IEA) | 1 | 9 | 0 |
| | Gramene63 (IEA only) | 282 | 147 | 0 |
| *Medicago truncatula* A17 | GOMAP | 2673 | 10 | 798 |
| | Gold Standard Gramene 63 (no IEA) | 2 | 23 | 0 |
| | Gramene63 (IEA only) | 309 | 243 | 0 |
| | Phytozome12 | 132 | 17 | 0 |
| *Medicago truncatula* R108 | GOMAP | 4168 | 7 | 803 |
| *Oryza sativa* | GOMAP | 1642 | 7 | 869 |
| | Gold Standard Gramene 63 (no IEA) | 37 | 833 | 0 |
| | Gramene63 (IEA only) | 238 | 64 | 0 |
| | Phytozome12 | 119 | 19 | 0 |
| *Phaseolus vulgaris* | GOMAP | 1190 | 6 | 783 |
| *Pinus lambertiana* | GOMAP | 1839 | 4 | 587 |
| *Sorghum bicolor* | GOMAP | 2384 | 66 | 783 |
| | Gold Standard Gramene 63 (no IEA) | 178 | 219 | 0 |
| | Gramene63 (IEA only) | 278 | 198 | 0 |
| | Phytozome12 | 131 | 12 | 0 |
| *Triticum aestivum* | GOMAP | 9624 | 17 | 1132 |
| | Gold Standard Gramene 63 (no IEA) | 1 | 5 | 0 |
| | Gramene63 (IEA only) | 584 | 319 | 0 |
| *Vigna unguiculata* | GOMAP | 1269 | 6 | 811 |
| | Phytozome12 | 122 | 27 | 0 |
| *Zea mays* B73.v4 | GOMAP | 2077 | 89 | 848 |
| | Gold Standard Gramene 63 (no IEA) | 50 | 633 | 0 |
| | Gramene63 (IEA only) | 306 | 140 | 0 |
| *Zea mays* Mo17 | GOMAP | 2346 | 83 | 823 |
| | Gold Standard Gramene 63 (no IEA) | 36 | 1489 | 0 |
| *Zea mays* PH207 | GOMAP | 2676 | 82 | 830 |
| | Gold Standard Gramene 63 (no IEA) | 37 | 2702 | 0 |
| *Zea mays* W22 | GOMAP | 2681 | 88 | 840 |
| | Gold Standard Gramene 63 (no IEA) | 30 | 499 | 0 |

Download this table (CSV)

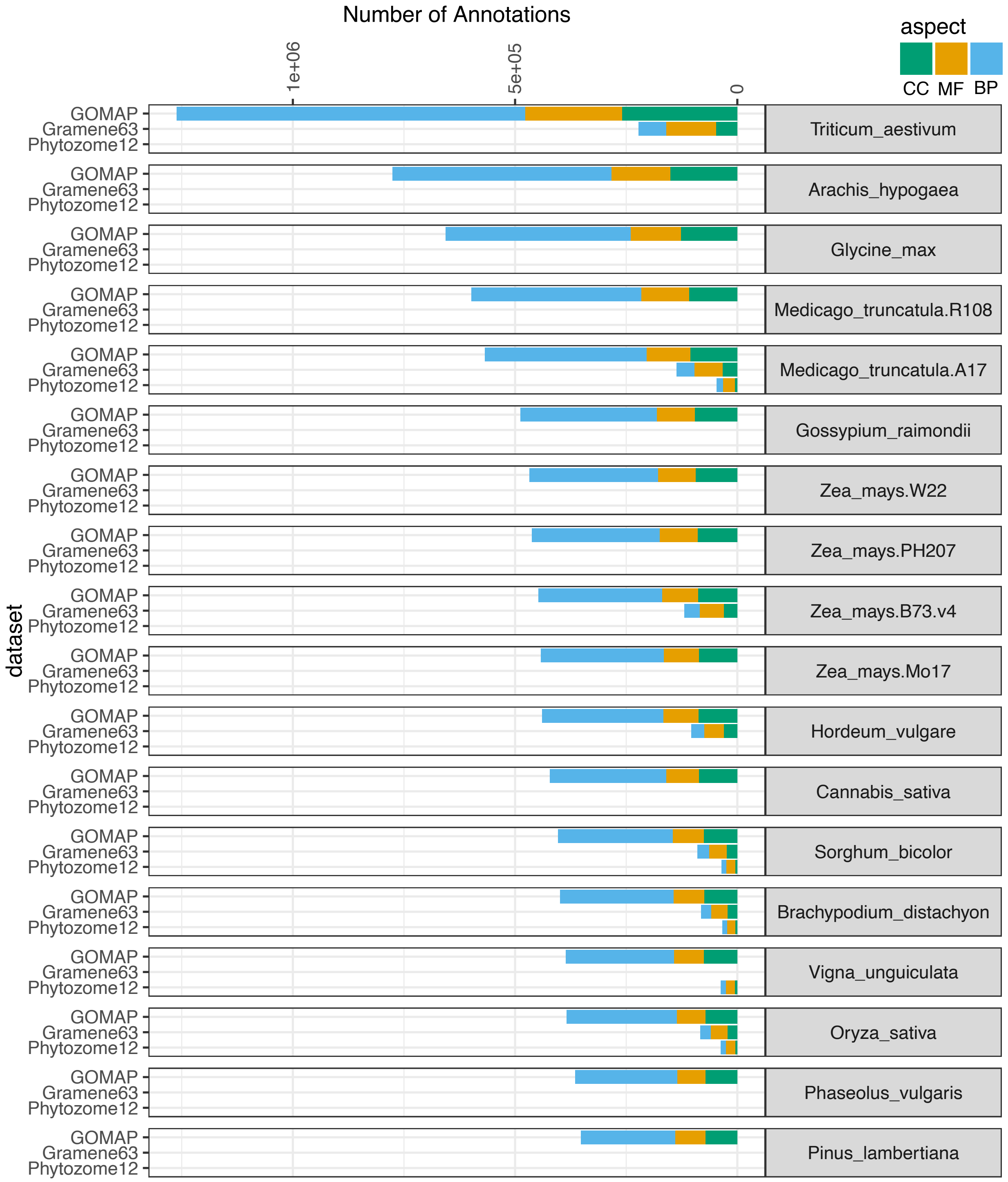


Supplementary Figure S1: Number of total annotations in each GO IEA dataset analyzed, colored by GO aspect (Cellular Component in green, Molecular Function in orange, and Biological Process in blue). Species are ordered based on the number of GO terms in the GOMAP dataset, with the species producing the most GO terms (*Triticum*) on top and the least (*Pinus*) on bottom.

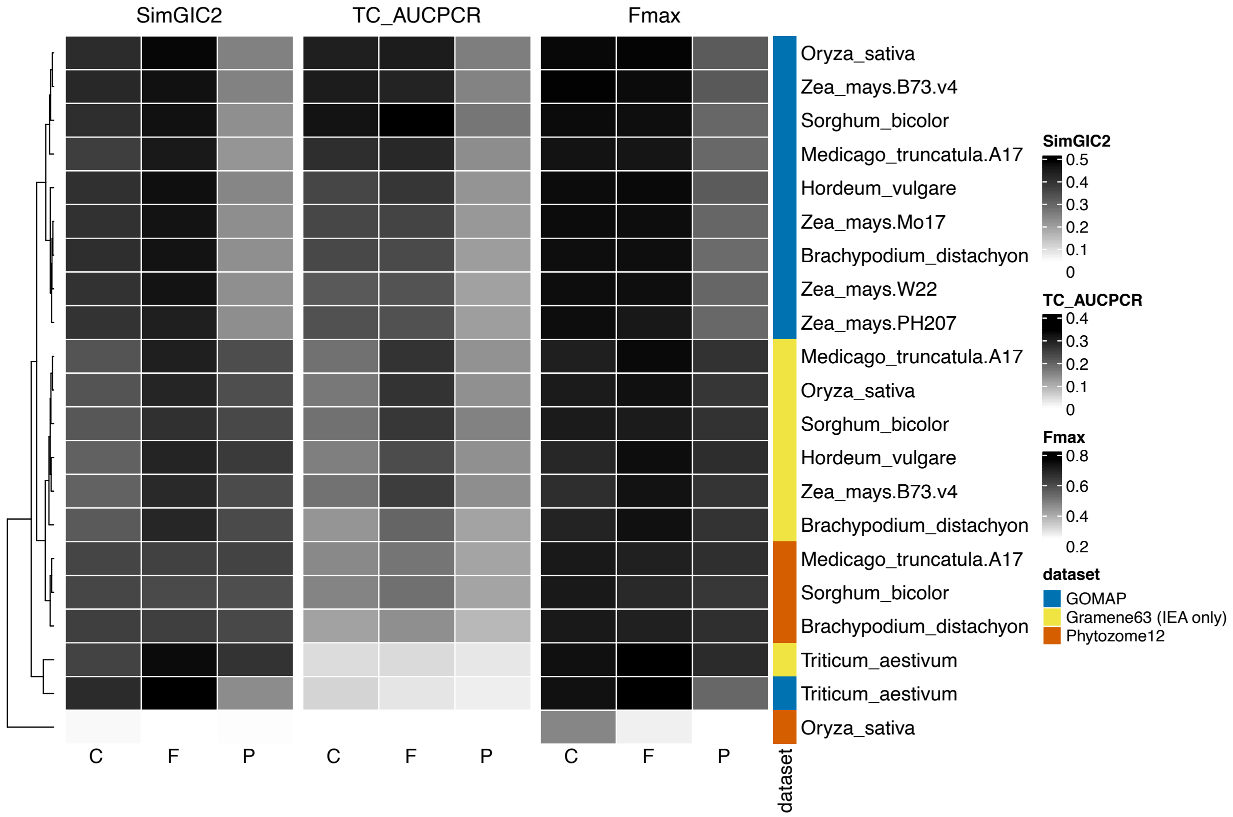


Supplementary Figure S2: Quality scores of the predicted annotation visualized as a grayscale heatmap. Each row represents a single species from a single data source. Input gene model dataset origin is indicated - blue for GOMAP, yellow for Gramene63, and orange for Phytozome12. Across the top are the three metric types used to assess the annotation quality. Across the bottom are subgraph indicators: C is Cellular Component, F is Molecular Function, and P is Biological Process. Darker cells indicate a higher (better) score whereas lighter cells indicate a lower score. Note that scales are different for each metric type (meaning that comparisons across the three metric types are not meaningful). Rows are clustered by pairwise correlation/similarity across all metrics. The dendrogram at left retraces clustering.

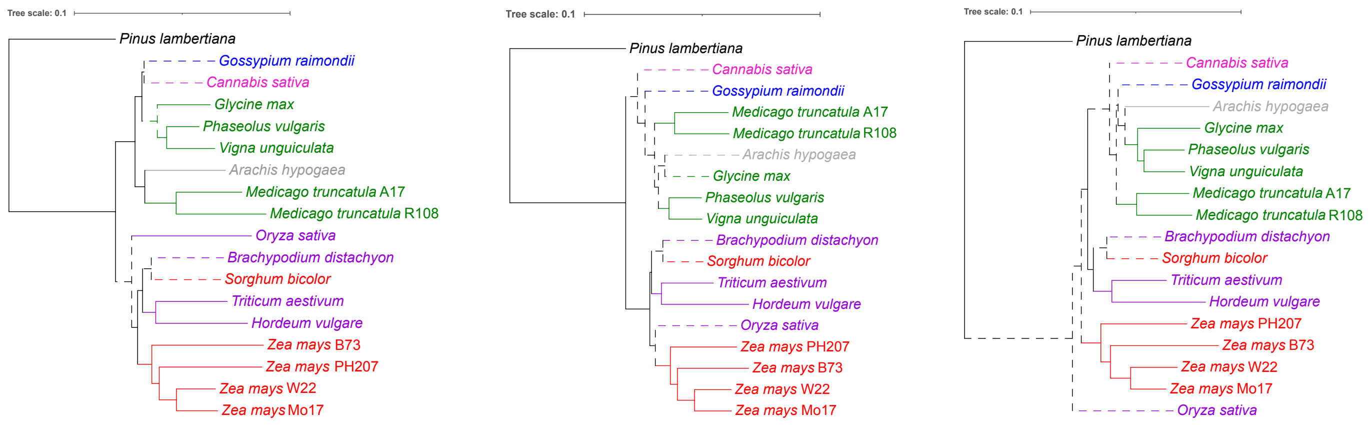


(a) Cellular Component (b) Molecular Function (c) Biological Process

Supplementary Figure S3: Neighbor-joining trees built on annotation subsets per GO aspect. Phylograms are colored and rooted as described in Figure 2. For each tree, only the annotations from the respective aspect of the Gene Ontology were used.
