## Supplementary figures and images for "Standardized genome-wide function prediction enables comparative functional genomics: a new application area for Gene Ontologies in plants"

### Supplemental Figure 1

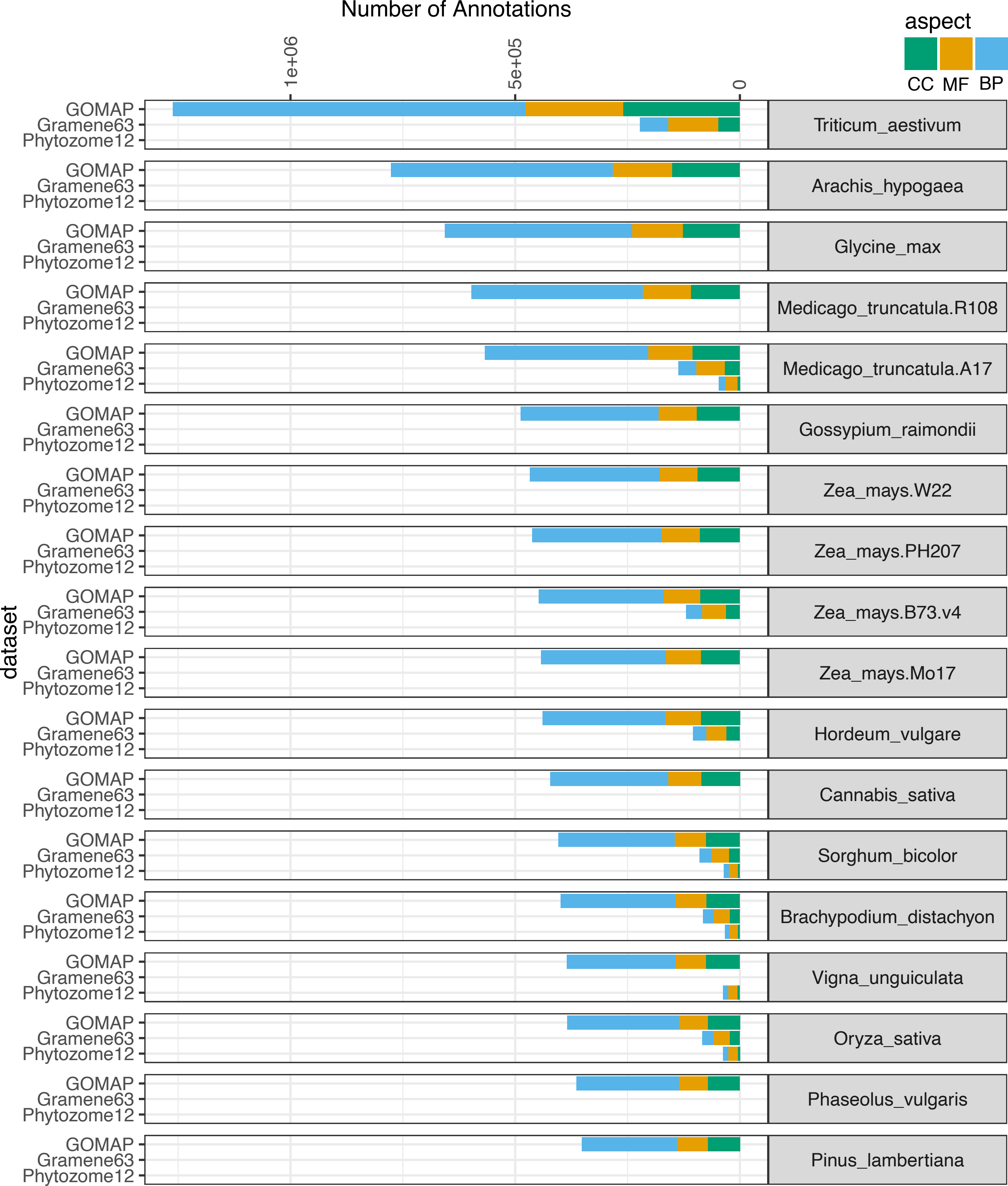

### Supplemental Figure 2

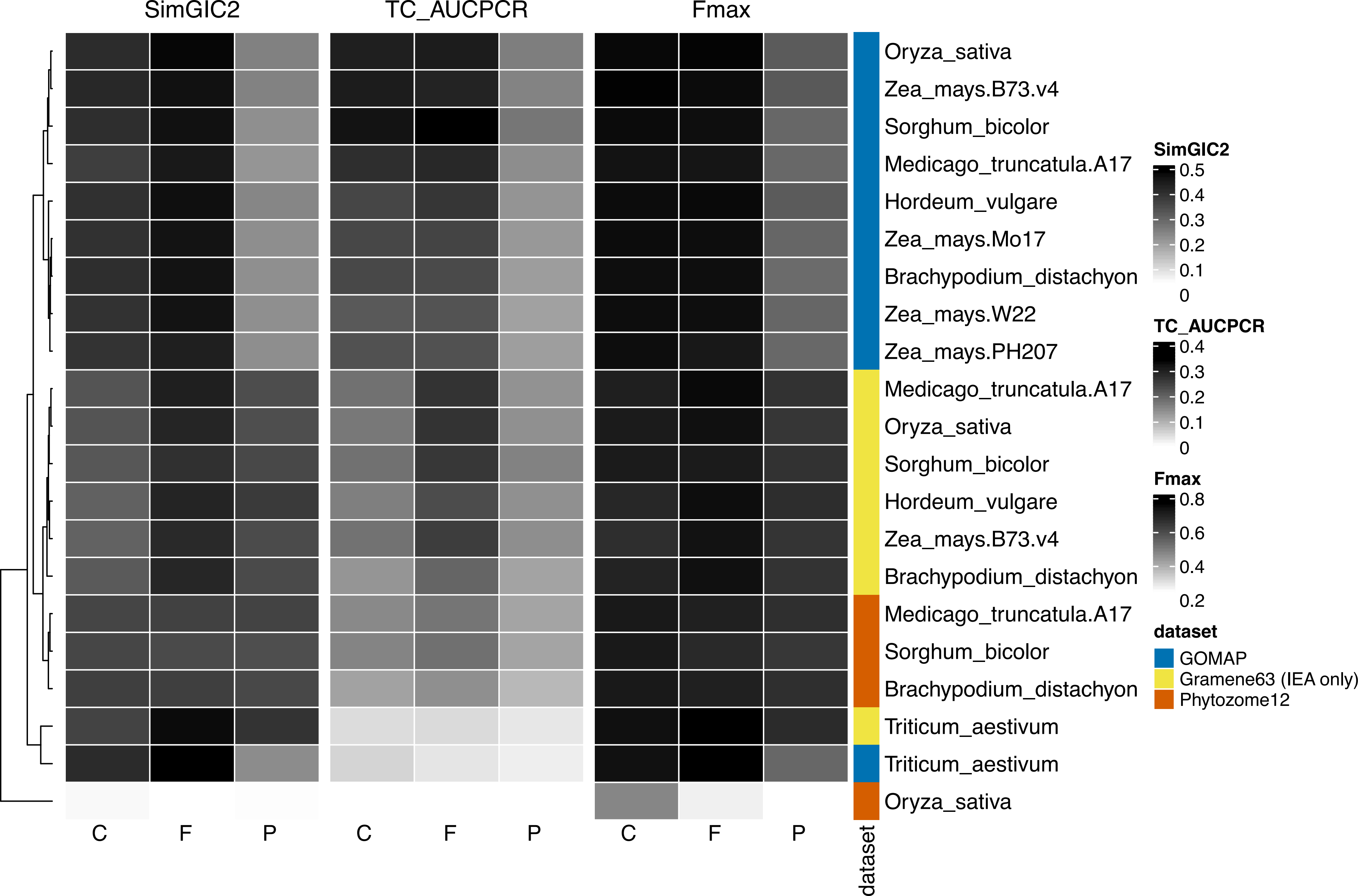

### Supplemental Figure 3a

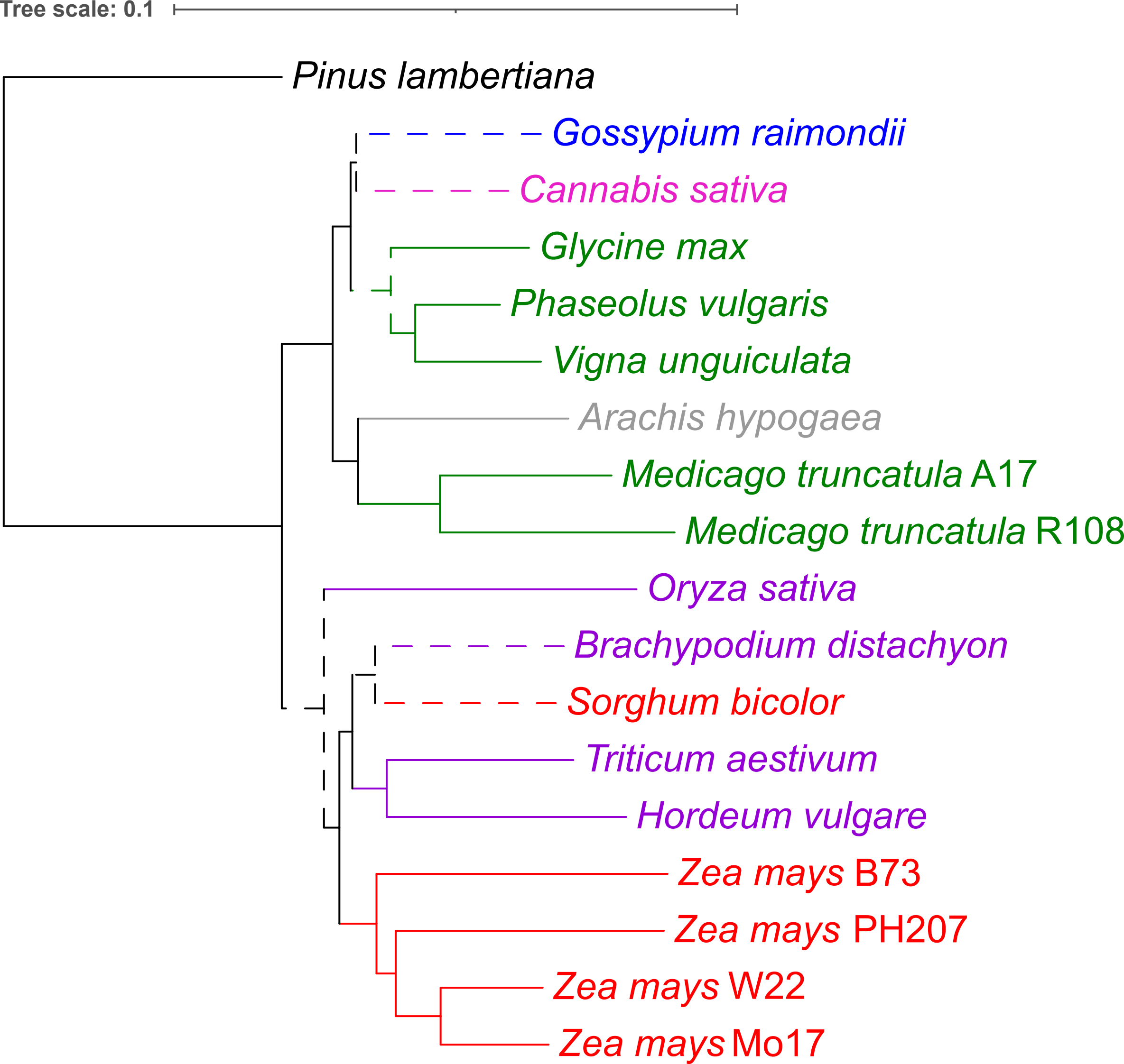

### Supplemental Figure 3b

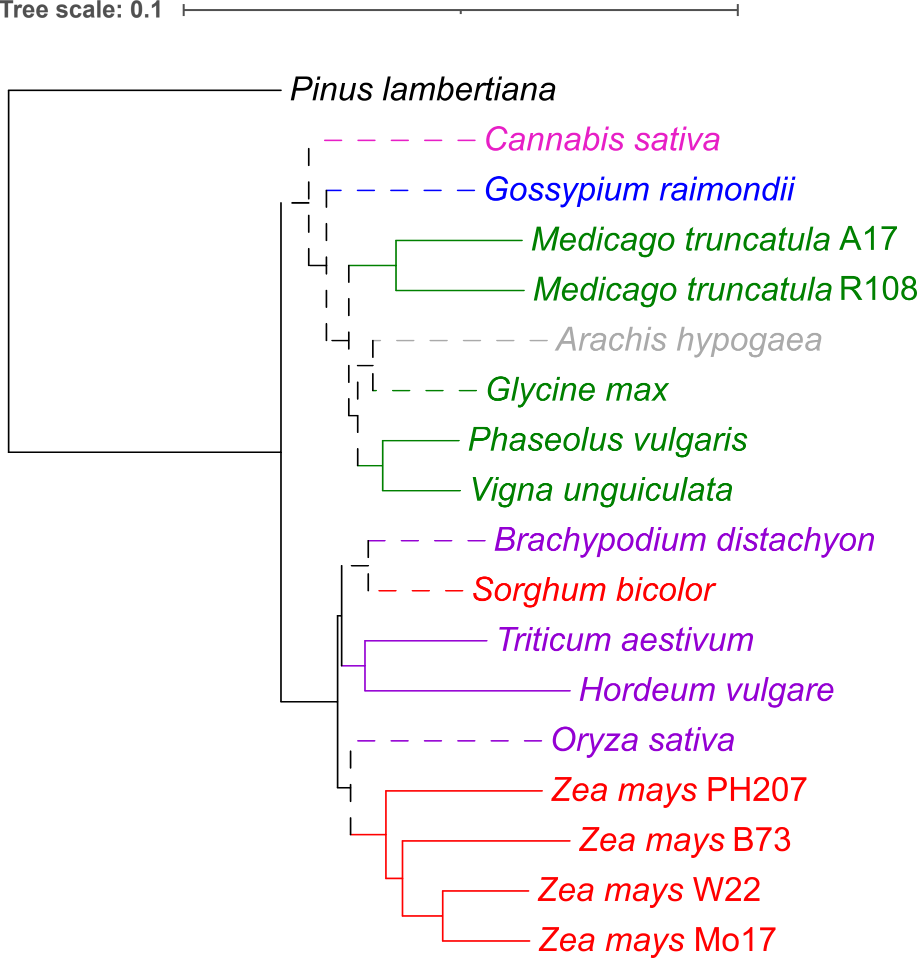

### Supplemental Figure 3c

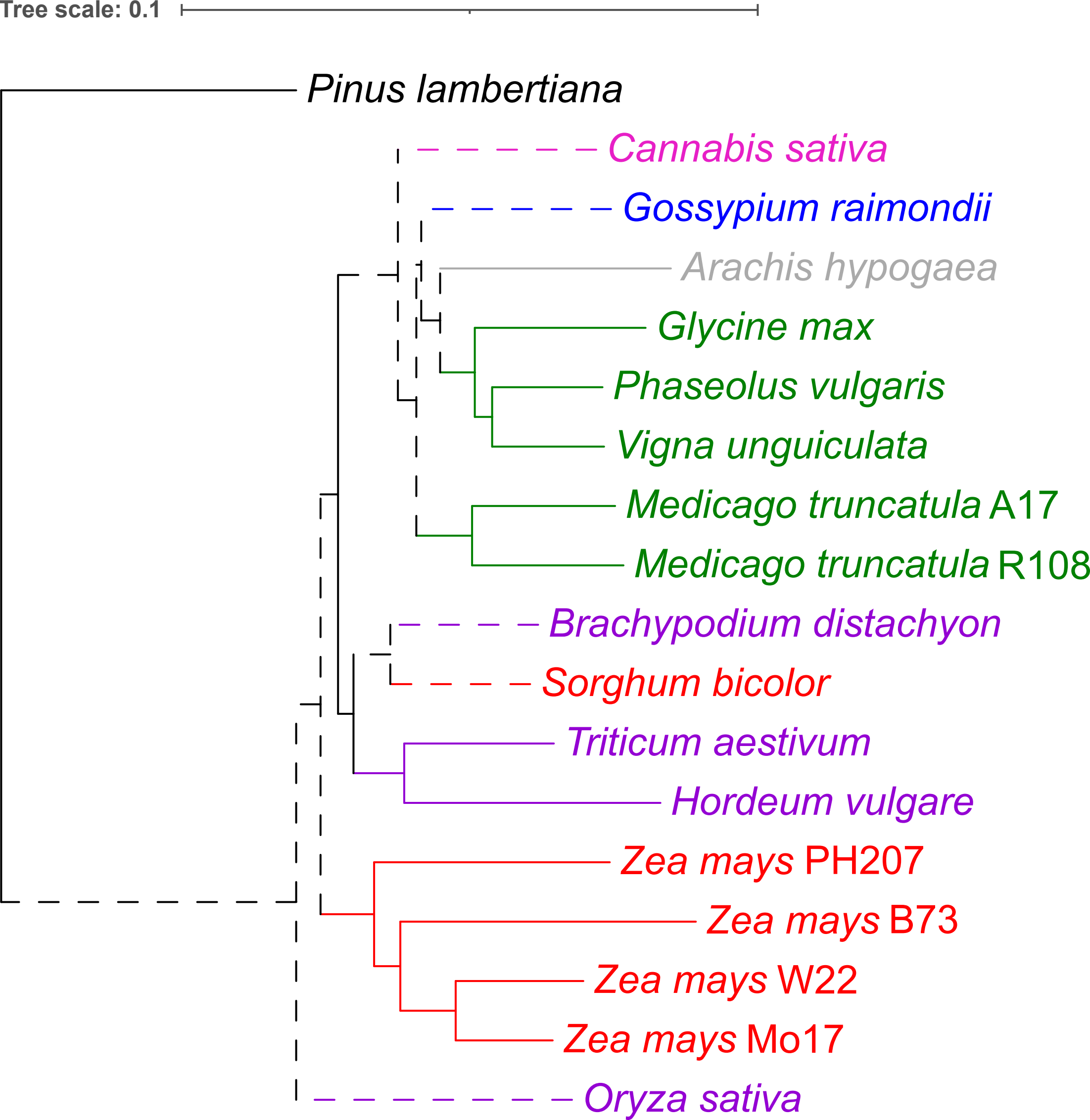
